# Beyond the Billion: Dose-Response Immunophenotyping and Machine Learning Classification of Live versus Heat-Treated Gram-Positive Probiotic Strains in Human Peripheral Blood Mononuclear Cells

**DOI:** 10.64898/2026.05.26.727599

**Authors:** Giovanni Deusebio, Annalisa Visciglia, Angela Amoruso, Marco Pane

## Abstract

Probiotic research is constrained by three pervasive yet insufficiently challenged assumptions: the requirement for a minimum of one billion colony-forming units for efficacy, the necessity for gut colonization, and the inherent superiority of live over heat-treated preparations. This study addresses these gaps through a fully factorial experimental design evaluating ten Gram-positive probiotic strains in both viable (Active Fluorescent Units, AFU) and heat-treated (Total Fluorescent Units, TFU) forms across three flow cytometry-verified concentrations (10^5^, 10^6^, 10^7^ cells/well per ISO 19344:2015) in primary human peripheral blood mononuclear cells (PBMC) from a single healthy male Caucasian donor (58 years), with simultaneous quantification of 17 cytokines by BioPlex suspension array. Viable preparations induced profoundly greater absolute cytokine responses than heat-treated preparations across 16 of 17 analytes, heat-treated preparations demonstrated stronger dose-response correlations (mean within-strain Spearman ρ up to 1.00) for 13 of 17 cytokines, a finding we attribute to the uncontrolled proliferation of live bacteria during 24-hour co-culture compressing the effective concentration range. Six of ten viable strains exhibited monotonically increasing profiles; two strains displayed non-monotonic bell-shaped kinetics with peak activity at 10^6^ AFU/well and significant attenuation at 10^7^, directly falsifying the assumption that dose escalation uniformly increases immunological activity. MCP-1 was the sole cytokine showing no significant difference between viability states at the observation level (p = 0.61, fold-change 1.0), providing an internal methodological control. In this single-donor model, unsupervised hierarchical clustering identified three immunological phenotype clusters, requiring multi-donor validation before these groupings can be treated as generalizable biological phenotypes, with Random Forest classification achieving 83.3% internal partition-recovery consistency (clusters derived from the same data; not an estimate of generalization to novel strains) versus 33.3% chance; In this single-donor experiment, IL-12p70, IL-13, and IFN-γ, not IL-6 or IL-1β, were the primary discriminators of strain identity; generalizability of this ranking requires multi-donor validation. Heat-treated preparations achieved ≥70% functional equivalence relative to viable preparations at 10^7^ TFU/well for the majority of responsive strains (Functional Equivalence Dose, FED^70^), while one strain showed strongly attenuated activity in heat-treated form across all concentrations, a finding subject to the caveat that no positive control stimulus was included to formally verify PBMC functional competence on the experimental day. These findings establish a methodological framework integrating flow cytometric standardization, multiplex immunophenotyping, and machine learning for evidence-based dose characterization, postbiotic functional equivalence assessment, and data-driven strain classification in probiotic research (all p-values are descriptive within a single-donor experimental context).

## 1. Introduction

The global probiotic market, valued at over 60 billion US dollars, rests on a scientific foundation that leading researchers increasingly regard as insufficient (Suez et al., 2019). Every probiotic product on the market declares a dose, yet none can provide rigorous evidence that the declared colony-forming unit count—whether one billion, ten billion, or two hundred billion—was selected based on biologically defined effective concentrations rather than manufacturing feasibility and competitive positioning. The International Scientific Association for Probiotics and Prebiotics defined probiotics as “live microorganisms that, when administered in adequate amounts, confer a health benefit on the host” (Hill et al., 2014), yet what constitutes “adequate amounts” remains remarkably poorly characterized at the strain level. The foundational FAO/WHO consensus encoded the phrase “adequate amounts” without any operational framework to define it (FAO/WHO, 2001), and the widespread assumption that a minimum of one billion colony-forming units (CFU) is required for efficacy has become an industry standard despite the absence of rigorous dose-ranging evidence for most strains and endpoints (Ouwehand, 2017). The regulatory consequences of this evidence vacuum are stark: the European Food Safety Authority has rejected every probiotic health claim submitted—over 100 applications—partly due to insufficient dose characterization (EFSA, 2012), the United States Food and Drug Administration considers probiotic disease claims as drug uses requiring full Investigational New Drug and New Drug Application pathways with no probiotic health claim approved to date, and the ICH E4 guideline mandates dose-response assessment as “an integral component of drug development” with a minimum of three dosage levels—a standard essentially absent in the probiotic field. Ouwehand (2017) reviewed seven meta-analyses spanning major probiotic endpoints and found no clear dose-response relationship for immune markers, bowel function, or general health outcomes. Only antibiotic-associated diarrhea showed a threshold effect near 10^10^ CFU per day. Even in vulnerable populations where dose optimization would carry the greatest clinical consequence, the evidence remains thin: Rath C et al. (2023) found limited dose-response evidence in preterm infants, with one randomized controlled trial showing no significant differences between high- and low-dose groups. The International Scientific Association for Probiotics and Prebiotics working group explicitly criticized the field, noting that “multiple doses are needed in order to elucidate the presence of dose-dependence” and that “single-dose studies are rarely acceptable” (Shane et al., 2010). The World Gastroenterology Organisation acknowledges that effective probiotic doses range from 10^8^ CFU per day for *Bifidobacterium longum* 35624 in irritable bowel syndrome to 450 billion CFU thrice daily for VSL#3—spanning a greater than 1,000-fold range—yet almost no studies have characterized the dose-response curve shape (WGO, 2023). In the only published five-dose-level human immune study, Christensen et al. (2006) found no significant immune changes at any dose level from 10⁸ to 10¹¹ CFU per day in 71 healthy adults, illustrating both the scarcity of dose-ranging studies and the challenge of translating in vitro findings to clinical immunological endpoints. Despite these admonitions, the vast majority of published probiotic studies employ a single dose without pharmacological justification, a practice that would be unacceptable in pharmaceutical development where ICH E4 guidelines mandate dose-response assessment with at least three dosage levels plus placebo. Health Canada takes a more progressive strain-specific approach but still requires evidence to support the daily dose for each strain—a requirement that presupposes the existence of dose-response data that, for immunological endpoints, the field has largely failed to generate. A second foundational assumption—that colonization of the gut is required for efficacy—has been systematically challenged. Zmora et al. (2018) used invasive endoscopic sampling to demonstrate that mucosal probiotic colonization is highly person-, region-, and strain-specific, and that stool analysis does not reflect mucosal colonization status. The companion study by Suez et al. (2018) showed that probiotic colonization after antibiotics actually delayed indigenous microbiome reconstitution versus spontaneous recovery or autologous fecal microbiota transplantation.

Kristensen et al. (2016) found no evidence that probiotics alter fecal microbiota composition in healthy adults across seven randomized controlled trials, suggesting that effects derive from transient metabolic activity and immune modulation rather than colonization. Suez et al. (2019) further highlighted that strain-level evidence is often inappropriately extrapolated across species, safety is under-evaluated, and personalized immune responses are routinely ignored. If colonization is neither reliable nor necessary, then the viability requirement embedded in the canonical definition itself demands re-examination.

This re-examination has gained formal recognition through the International Scientific Association for Probiotics and Prebiotics definition of postbiotics as “preparations of inanimate microorganisms and/or their components that confer a health benefit on the host” (Salminen et al., 2021), which explicitly requires deliberate inactivation, a defined source microorganism, and demonstrated health benefit. The clinical evidence supporting this paradigm has grown substantially. Depommier et al. (2019) demonstrated in a randomized, double-blind, placebo-controlled trial that pasteurized *Akkermansia muciniphila* significantly improved insulin sensitivity, reduced insulinemia, and lowered total cholesterol, while the live form did not reach significance on primary endpoints. A systematic review of 40 randomized controlled trials encompassing 3,913 participants found that modified probiotics were not significantly different from living probiotics in 86% of preventive and 69% of treatment trials, with treated bacteria actually outperforming live preparations in 15% of treatment trials (Zorzela et al., 2017). The largest randomized controlled trial of a heat-treated bacterium for irritable bowel syndrome further reinforced this evidence: Andresen et al. (2020) demonstrated that heat-treated *Bifidobacterium bifidum* MIMBb75 achieved a 34% versus 19% placebo response rate in 443 patients. These findings raise a fundamental question: if heat-treated bacteria can match or exceed the efficacy of live preparations, then the requirement for viability—and the entire construct of “adequate amounts” based on CFU enumeration of viable cells—requires reconsideration.

This reconsideration is further complicated by the quantification crisis pervading probiotic research. Colony-forming unit plate counting, the industry standard, systematically underestimates viable cell populations by missing viable-but-non-culturable cells (Davis, 2014). A landmark interlaboratory study across 244 laboratories demonstrated that optical density at 600 nm calibrated against CFU produced values more than an order of magnitude different from flow cytometry ground truth, with the same optical density reading yielding CFU estimates spanning a 3.3-fold range between instruments (Beal et al., 2020). Flow cytometry enumeration has been shown to detect approximately 70% more viable cells than plate counting for the same product (Visciglia et al., 2022), with precision nearly twice that of conventional methods: relative standard deviation of 6.70% versus 12.52% for plate counting (Tracey et al., 2023). Reproducibility between standardized plate count methods ranges from 0.5 to 1 log, meaning results can differ up to 10-fold between laboratories (Foglia et al., 2020). The discrepancy widens with storage time as cells transition to the viable-but-non-culturable state: Gorsuch et al. (2019) showed that flow cytometric and quantitative polymerase chain reaction counts were similar and substantially higher than plate counts at later storage times, rendering CFU-based dosing increasingly unreliable for aged preparations. Critically, CFU methodology is entirely inapplicable to heat-treated preparations by definition. A 2024 Frontiers in Microbiology study (Bolzon et al., 2024) developed the first optimized flow cytometric method for tyndallized bacterial cells using TO/PI double staining. The regulatory adoption of flow cytometric total fluorescent unit enumeration for pasteurized *Akkermansia muciniphila*—the first next-generation bacterium to receive EFSA novel food approval (EFSA, 2021)—established the regulatory precedent for culture-independent quantification. Flow cytometry enumeration, standardized through ISO 19344 and validated across 15 laboratories in 5 countries, provides the methodological foundation for quantifying both viable and heat-treated preparations with equivalent accuracy capability that proved decisive for the present study’s direct comparison of live and heat-treated bacteria at matched concentrations.

### 1.1 Cell Model: Rationale, Validation, and Analytical Gaps

The rationale for the PBMC single-donor model, its analytical validation, and the specific analytical gaps addressed by this study are detailed in the Supplementary Material (Section S1).

## 2. Materials and Methods

### 2.1 Study Design and Experimental Overview

This study employed a fully factorial experimental design to evaluate the immunomodulatory properties of ten distinct Gram-positive probiotic bacterial strains on human peripheral blood mononuclear cells. The experimental factors comprised: bacterial strain identity (10 levels, designated A through L, excluding J and K); bacterial viability status (2 levels: live and heat-treated); and bacterial concentration (3 levels: 10^5, 10^6, and 10^7 AFU or TFU/mL). Each experimental condition was assayed in technical triplicate, yielding a total of 180 treatment observations. Unstimulated peripheral blood mononuclear cell cultures served as negative controls (12 replicate wells); cytokine concentrations from the 12 wells were averaged per analyte prior to fold-change calculation, yielding a single negative control reference mean per cytokine used as the denominator for all conditions. The primary endpoint was the quantification of 17 cytokines in culture supernatants following 24-hour co-incubation.

### 2.2 PBMC Isolation and Culture

Cryopreserved peripheral blood mononuclear cells were obtained from Lonza (Basel, Switzerland) as a pre-characterized preparation from a single healthy male Caucasian donor (58 years of age). Cells were thawed according to the manufacturer’s protocol, washed twice with pre-warmed complete RPMI-1640 medium, and resuspended in complete medium supplemented with 10% heat-inactivated fetal bovine serum (FBS; Thermo Fisher Scientific, Waltham, MA, USA) and 2 mM L-glutamine.

### 2.3 Probiotic Strain Preparation

Ten probiotic strains (designated A–L for this analysis; full taxonomic characterization will be disclosed in subsequent publications) were maintained as cryostocks and revived by sequential subculturing in appropriate liquid growth media. All strains were used at the second passage post-thaw to ensure consistent physiological state. For stimulation suspensions, strains were harvested from fresh broth culture at late-exponential phase by centrifugation, washed once, and resuspended in sterile deionized water to eliminate potential confounding contributions of buffer components to PBMC cytokine responses.

#### 2.3.1 Thermal Treatment Protocol

Heat-treated (HT) preparations were produced from the same washed bacterial suspensions as viable preparations. A standardized thermal treatment protocol was applied identically across all ten strains: exposure to temperatures exceeding 85°C for 60 minutes. Complete treatment was confirmed by plating on appropriate solid recovery medium and confirming absence of colony growth after 48–72 hours at optimal temperature for every batch and strain.

#### 2.3.2 Flow Cytometric Enumeration: AFU and TFU Standardization

Quantification of both viable and heat-treated suspensions was performed by flow cytometry per ISO 19344:2015 using a NovoCyte flow cytometer (Agilent Technologies, Santa Clara, CA, USA) and Thiazole Orange (TO)/Propidium Iodide (PI) dual-fluorochrome staining. TO labels all bacteria regardless of membrane integrity; PI selectively penetrates membrane-compromised cells. Viable bacteria (TO^+^/PI^−^) were quantified as Active Fluorescent Units per well (AFU/well); heat-treated bacteria (TO^+^/PI^+^) as Total Fluorescent Units per well (TFU/well). This approach, validated for heat-treated bacterial cells using TO/PI staining (PMC11099204, 2024), enables direct, concentration-matched comparison between viable and heat-treated preparations in identical physical units (cells per well), independently of culturability — resolving the fundamental CFU limitation identified by Davis (2014), Beal et al. (2020), and Foglia et al. (2020). Final concentrations of 10^5^, 10^6^, and 10^7^ AFU/well or TFU/well were confirmed by post-dilution re-measurement prior to addition to PBMC cultures.

### 2.4 PBMC-Bacteria Co-culture

Peripheral blood mononuclear cells were seeded in 48-well flat-bottom plates at a density of 1 × 10^6 cells per well in 1ml antibiotic-free RPMI-1640 medium supplemented with 10% heat-inactivated fetal bovine serum and 2 mM L-glutamine. Bacterial preparations (live or heat-treated) were added at the designated concentrations. Co-cultures were incubated at 37°C in a humidified atmosphere containing 5% CO₂ for 24 hours. Following incubation, culture plates were centrifuged at 300 × g for 10 minutes, and cell-free supernatants were carefully collected and stored at −80°C until cytokine quantification.

### 2.5 Multiplex Cytokine Quantification

Cytokine concentrations in culture supernatants were quantified using the Bio-Plex Pro Human Cytokine 17-plex Assay (Bio-Rad Laboratories, Hercules, CA, USA) according to the manufacturer’s instructions. The panel comprised: G-CSF, GM-CSF, IFN-γ, IL-1β, IL-2, IL-4, IL-5, IL-6, IL-7, IL-8/CXCL8, IL-10, IL-12p70, IL-13, IL-17A, MCP-1/CCL2, MIP-1β/CCL4, and TNF-α. Eight-point standard curves were generated for each cytokine, and concentrations were calculated by 5-parameter logistic regression curve fitting. Quality control criteria included intra-assay CV ≤ 15% and inter-assay CV ≤ 20%.

### 2.6 Data Analysis and Statistics

Immunomodulatory activity was expressed as fold-change (FC) relative to mean cytokine concentration in unstimulated control wells of the same experimental run. Data are presented as mean ± SD across three technical replicates. The negative control denominator represents the intra-analyte mean of 12 replicate wells. Consequently, fold-change SD values reported for each condition reflect within-condition technical variability only, and do not capture uncertainty in the negative control reference value.

#### 2.6.1 Data Preprocessing

Out-of-range values below the detection limit were replaced with the minimum concentration observed for that analyte across all experimental samples in the dataset. Out-of-range values exceeding the upper quantification limit were replaced with the maximum concentration observed for that analyte across all experimental samples in the dataset. This approach anchors substituted values to the empirically determined quantification boundaries of the specific assay run, avoiding arbitrary scalar multipliers. Log^10^-transformation was applied to normalize distributions and stabilize variance. For multivariate analyses, RobustScaler normalization (using median and interquartile range) was employed to reduce the influence of extreme outliers.

#### 2.6.2 Differential Expression and Dose-Response Analysis

Differences in cytokine expression between viable and heat-treated conditions were evaluated using Mann-Whitney U tests with Benjamini-Hochberg false discovery rate correction (Benjamini and Hochberg, 1995). It is noted that the 30 condition means per viability group are not fully independent observations, as they derive from a single biological donor with n = 3 technical replicates per condition. This pseudoreplication architecture means that the Mann-Whitney U p-values reported throughout this study should be interpreted as descriptive indicators of effect magnitude and direction rather than as inferential probability statements generalizable to a broader population. All statistical comparisons are reported in this spirit — as structured descriptions of the observed data within this single-donor experiment, consistent with the exploratory and hypothesis-generating design of the study. Fold-change was calculated as the ratio of geometric means. Dose-response relationships were assessed using Spearman rank correlation between bacterial concentration and cytokine levels, stratified by viability status. Kruskal-Wallis tests were used to evaluate overall concentration effects within each viability state.

#### 2.6.3 Dimensionality Reduction and Clustering

Principal component analysis was performed on RobustScaler-standardized log-transformed data. Component retention was determined using the Kaiser criterion and scree plot inspection, targeting at least 80% cumulative variance explained. K-means clustering was applied to the full dataset with the optimal cluster number determined by maximizing both silhouette score and Calinski-Harabasz index across k = 2 to 10. Hierarchical clustering using Ward’s minimum variance method with Euclidean distance provided complementary validation of the partition structure. PCA was applied at two levels: (1) on the full observation matrix (192 observations × 17 cytokines) to characterize the global variance structure of the dataset including technical replicates, and (2) on the condition-mean matrix (60 conditions × 14 cytokines, means of technical replicates) to visualize the immunological space occupied by each strain × viability × dose combination and support hierarchical clustering of strain-level fingerprints.

#### 2.6.4 Software

All analyses were performed in Python 3.12 using scikit-learn (Pedregosa et al., 2011), scipy, pandas, xgboost, matplotlib, and seaborn. Complete analysis scripts will be made available upon acceptance.

##### AI Disclosure

Generative artificial intelligence (Claude, Anthropic, claude.ai) was used in the preparation of this manuscript for scientific writing and structural editing, Python scripting for figure generation and machine learning analysis, and literature synthesis. All AI-generated content was critically reviewed, verified against primary sources, and edited by the authors. All interpretations, conclusions, and scientific claims are the sole responsibility of the authors. This disclosure is made in accordance with Frontiers in Microbiology editorial policy.

#### 2.6.5 Power Considerations

Formal prospective power analysis is not directly applicable to a single-donor co-culture design with n = 3 technical replicates per condition, as the biological unit of observation is the condition mean rather than an independent donor. The design was sized to enable comprehensive factorial coverage (10 strains × 2 viability states × 3 concentrations × 3 technical replicates) within a single controlled experiment rather than to achieve a pre-specified statistical power for a primary endpoint. The primary viability contrast (30 condition means per group) is sufficiently powered to detect the large effect sizes observed (356.9-fold TNF-α, 72.6-fold MIP-1β) by orders of magnitude. For smaller effects — including strain × dose interactions and FED⁷⁰ convergence ratios — the three-replicate design provides descriptive precision rather than inferential power, consistent with the exploratory and hypothesis-generating framing of this study. Multi-donor sample size: based on the inter-donor cytokine production variability reported by Li et al. (2016) (CV 20–35% for stimulated PBMC), a minimum of 5–8 biological donors would be required to detect strain × dose interaction effects of 1.5-fold magnitude at 80% power (α = 0.05). This estimate informs the multi-donor validation plan described in Section 4.8

### 2.7 Machine Learning Analysis

#### 2.7.1 Classification Models

Four algorithms were evaluated for binary classification of bacterial viability status: Logistic Regression with elastic net regularization, Random Forest, Extreme Gradient Boosting (XGBoost; Wandong, 2022 and Guestrin, 2016), and Support Vector Machine with radial basis function kernel (Cortes and Vapnik, 1995). Hyperparameters were optimized via exhaustive grid search with 5-fold stratified cross-validation, using AUC-ROC as the primary optimization criterion. Pairwise model comparisons were performed using McNemar’s test. Feature importance was derived from model coefficients (Logistic Regression), Gini importance (Random Forest, XGBoost), and permutation importance (Support Vector Machine). All stochastic procedures used a fixed random seed (seed = 42) for reproducibility.

#### 2.7.2 Strain Classification and Functional Annotation

Random Forest classification (n_estimators = 500, class_weight =’balanced’, max_features =’sqrt’) was additionally applied to: (a) classify strains from viable preparations and compute cytokine feature importance by mean decrease in impurity; and (b) classify functional clusters from hierarchical clustering. All classification performance was evaluated by Leave-One-Out cross-validation (LOO-CV) and compared against the appropriate chance level (10% for 10-class strain, 33.3% for 3-cluster). Feature importance values in Figure 9B are reported as mean ± SD across 30 independent model fits generated by varying the random seed (seeds 1–30), to characterize the stability of importance rankings across stochastic variation in tree construction. Strain-specific immunomodulatory profiles were characterized using z-score standardization and functional composite scores for five predefined cytokine groups: pro-inflammatory (TNF-α, IL-1β, IL-6, IFN-γ), anti-inflammatory (IL-10, IL-4), Th1-associated (IFN-γ, IL-2, IL-12p70), Th2-associated (IL-4, IL-5, IL-13), and chemoattractant (IL-8, MCP-1, MIP-1β).

#### 2.7.3 Bayesian Estimation of Immunostimulatory Probability

The probability of each experimental condition producing immunostimulatory responses was estimated using a Bayesian framework with Jeffrey’s non-informative prior (Beta(0.5, 0.5)). For each strain × viability × concentration combination, a cytokine was classified as “elevated” if its mean concentration exceeded 150% of the negative control median. The 150% threshold was selected as a conservative criterion representing a 1.5-fold elevation above baseline, consistent with the minimum biologically meaningful fold-change in PBMC immunoassay systems given typical intra-assay variability of 15–20%. Jeffrey’s non-informative prior (Beta(0.5, 0.5)) was selected to minimize prior influence on posterior estimates; sensitivity analyses using a uniform prior (Beta(1, 1)) and a more conservative threshold (200% of control median) yielded qualitatively equivalent results, with all live conditions maintaining P(immunostimulatory) > 0.80 and the relative ordering of heat-treated conditions preserved. The number of elevated cytokines (out of 17) was used as the success count in a binomial model, yielding posterior Beta distributions for the probability of immunostimulation. It is acknowledged that the binomial model assumes independence between cytokine responses, an assumption that is not fully met given the high inter-cytokine correlations observed in this dataset (Spearman ρ up to 0.97; Section 3.8). The resulting posterior distributions should therefore be interpreted as descriptive summaries of response breadth rather than as independent probabilistic confirmation of the frequentist findings.

## 3. Results

### 3.1 Data Quality and Distribution Characteristics

The genome-wide immunomodulatory overview is shown in Figure 1A (heatmap) and Figure 1B (pathway-level radar). Data quality metrics, out-of-range handling and imputation, and distributional characteristics (Shapiro–Wilk normality testing) are reported in the Supplementary Material (Section S2); all downstream analyses met the stated quality criteria.

**Figure 1A.**
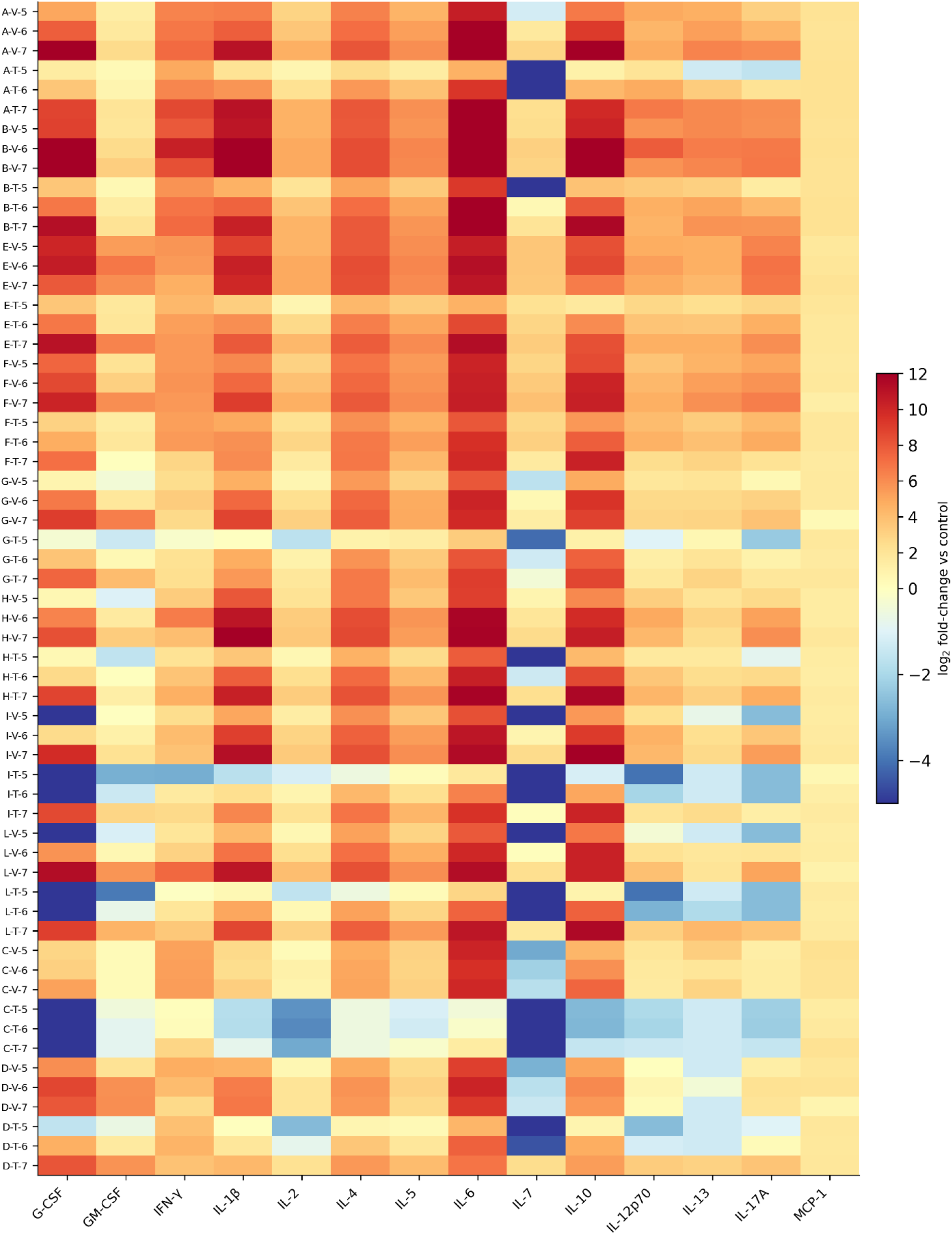
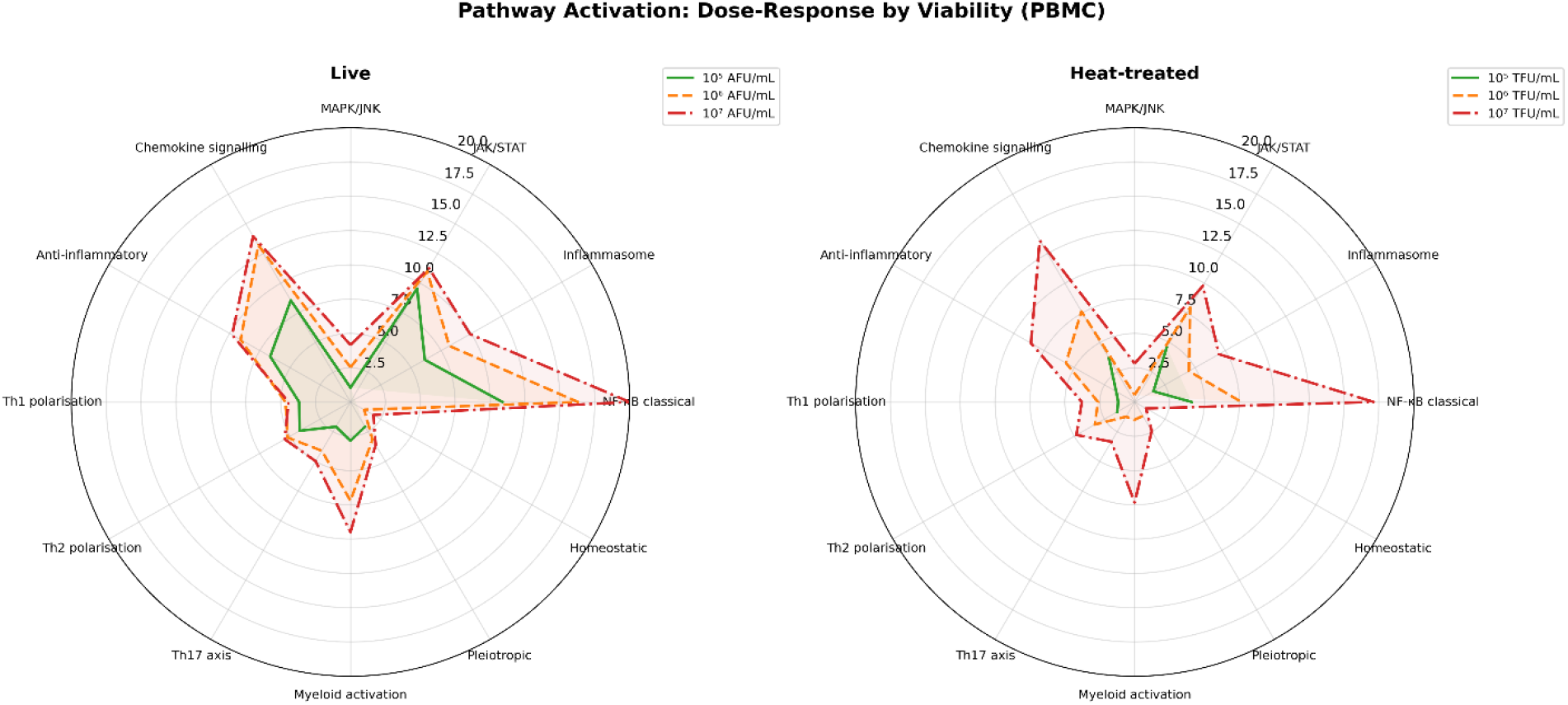
Overview of immunomodulatory activity across all strains, doses and preparation types. Heatmap of mean log₂ fold-change (n = 3 technical replicates) for 14 quantifiable cytokines relative to the unstimulated PBMC control (IL-8, MIP-1β, TNF-α excluded; see Supplementary Figure 1). Rows: strain (A–L), dose (10⁵, 10⁶, 10⁷ cells/well), preparation type (V = viable/AFU; HT = heat-treated/TFU). Colour: diverging RdYlBu, capped at log₂ = −5 (lower) and +12 (upper). White lines delineate strains. **Figure 1B.** Pathway-level radar visualisation of the same dataset. Mean immunomodulatory activity is summarised across twelve functional pathway categories for viable (left) and heat-treated (right) preparations, averaged across all ten strains at each of the three dose levels (10⁵ green solid, 10⁶ orange dashed, 10⁷ red dash-dotted). Each spoke represents one functional grouping of the 17-plex BioPlex panel: MAPK/JNK, JAK/STAT, inflammasome, NF-κB classical, homeostatic, pleiotropic, myeloid activation, Th17 axis, Th2 polarisation, Th1 polarisation, anti-inflammatory, and chemokine signalling. Independent radial scales are used for the two panels to accommodate the higher absolute activation magnitude of viable preparations. The concentric expansion of the polygons with increasing dose visualises the monotonic dose-dependent intensification of pathway engagement common to both preparation types. Anti-inflammatory, chemokine and inflammasome modules emerge as the dominant axes of response, while homeostatic and Th1-polarising modules show comparatively limited engagement. Pathway-category groupings represent a functional aggregation for visual integration; quantitative pathway-level inference is not derived from these aggregations.

### 3.2 Live Bacteria Induce Profoundly Greater Cytokine Responses Than Heat-treated Preparations

TNF-α showed the largest fold-change (356.9-fold higher in viable conditions), followed by G-CSF (15.4-fold), IL-1β (13.8-fold), IL-10 (9.9-fold), IL-6 (9.3-fold), IL-8 (8.9-fold), MIP-1β (72.6-fold), and IL-4 (5.5-fold) (Supplementary Figure 2). Pro-inflammatory innate cytokines were disproportionately affected: TNF-α, IL-1β, and IL-6 collectively demonstrated fold-changes exceeding 9-fold, consistent with potent NF-κB-dependent transcriptional activation by metabolically active bacteria. Among adaptive cytokines, IFN-γ (3.0-fold), IL-2 (3.8-fold), and IL-12p70 (4.0-fold) were significantly elevated in viable conditions, indicating Th1 activation. IL-4 (5.5-fold), IL-5 (3.1-fold), and IL-13 (2.9-fold) were also significantly higher, suggesting parallel Th2 activation rather than selective polarization. IL-17A (5.4-fold) showed substantial differences. IL-7 and IL-13 did not reach statistical significance after correction for multiple comparisons when analysed at the condition mean level (FDR p = 0.75 and p = 0.84, respectively), reflecting high inter-strain variability relative to the viability effect for these analytes, reflecting the high proportion of heat-treated conditions at the LLOQ floor for these analytes. MCP-1 (CCL2) was the sole cytokine showing no significant difference between viability states at the observation level (Mann-Whitney U p = 0.61; fold-change 1.0). All p-values reported in this section are descriptive indicators within a single-donor experiment and should not be interpreted as inferential estimates of population-level effects. This exception is biologically informative: MCP-1 is a potent monocyte chemoattractant produced through both MyD88-dependent and MyD88-independent signaling pathways, and its constitutive-like expression in both conditions suggests that the mere physical presence of bacterial structural components — regardless of viability — provides a sufficient stimulus for MCP-1 secretion. The viability-independence of MCP-1 across all ten strains and three concentrations constitutes an internal methodological control, confirming that the differential responses observed for the other 14 cytokines reflect genuine biological discrimination rather than experimental confounding.

All 14 significantly different cytokines exceeded both the statistical threshold (FDR < 0.05) and the magnitude threshold (absolute log₂ fold-change > 0.5), indicating biologically meaningful as well as statistically significant effects. The universal positive directionality — all 16 significant cytokines showing higher expression in viable conditions — demonstrates that viability universally enhances rather than selectively modulates the cytokine response, consistent with a generalized amplification mechanism rather than qualitative immune reprogramming.

#### 3.2.1 Bayesian Confirmation of Viability Effect

Bayesian estimation independently confirmed these findings: live bacteria conditions achieved P(immunostimulatory) ranging from 0.64 (Strain L at 10⁵ AFU/well) to 0.97, with 16 of 30 conditions reaching the maximum posterior probability of 0.97 (17/17 cytokines elevated above 150% of control median). In contrast, heat-treated conditions ranged from P = 0.19 (Strains C and I at 10⁵ TFU/well) to P = 0.97 (Strains E and F already at 10⁵ TFU/well; Strains B, D, H, L at 10⁷ TFU/well), demonstrating that the immunostimulatory gap between viability states is dose-dependent and strain-modulated (Figure 2, Figure 3).

**Figure 2.**
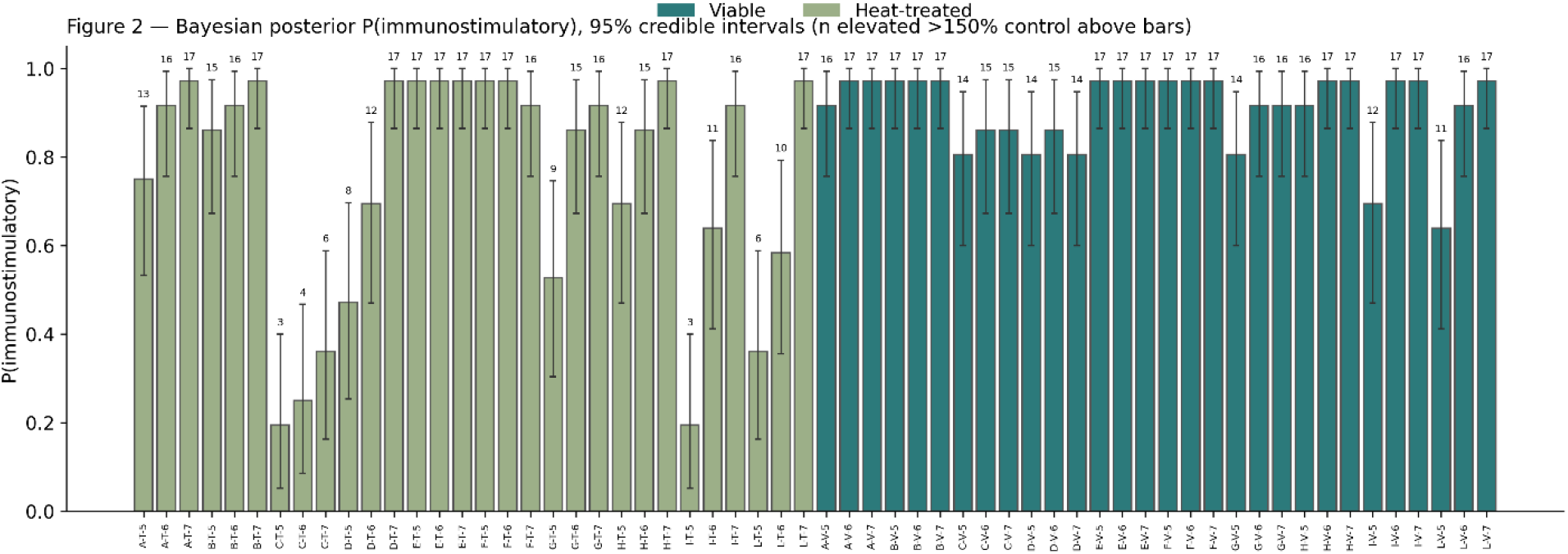
Bayesian posterior P(immunostimulatory) -95% credible intervals-for all conditions (number above bars: n cytokines elevated above 150% control)

**Figure 3.**
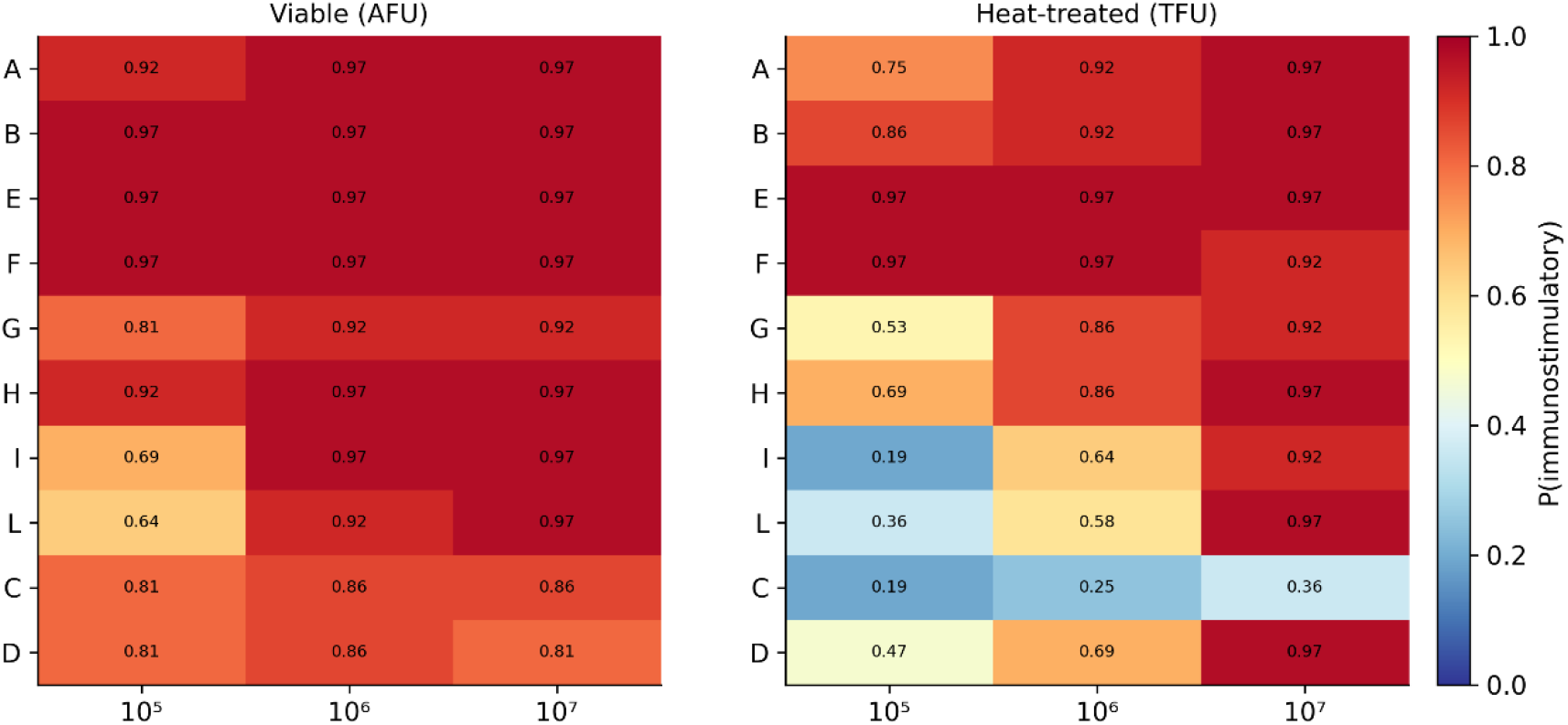
P(immunostimulatory) – strain x dose matrix

### 3.3 Dose-Response Relationships Differ Qualitatively Between Viability States

The second major finding — addressing whether measurable dose-response relationships exist — revealed an unexpected and methodologically significant asymmetry. Heat-treated preparations demonstrated stronger dose-response correlations overall: the strongest mean within-strain Spearman correlations were observed for G-CSF (mean ρ = 1.00), IL-4 (mean ρ = 1.00), MIP-1β (mean ρ = 1.00), IL-1β (mean ρ = 0.95), IL-10 (mean ρ = 0.95), and IL-6 (mean ρ = 0.95) in the heat-treated conditions. Given that Spearman rank correlation with three dose levels can only assume discrete values (0, ±0.5, ±1.0), formal p-value testing is not appropriate; direction and magnitude are reported descriptively. Heat-treated preparations showed stronger dose-response correlations than viable preparations for **13 of 17** cytokines when assessed as mean within-strain Spearman rank correlations across the three dose levels. Viable preparations, despite inducing substantially greater absolute cytokine levels at every concentration, exhibited weaker dose-response correlations across most analytes.

We hypothesize that live bacteria proliferate during incubation: even the lowest inoculum (10⁵ AFU/well) may have expanded substantially during the experiment, compressing the effective difference between the three nominal dose levels. This hypothesis is consistent with the observed asymmetry but was not directly tested in this study, as endpoint bacterial enumeration was not performed. Live bacteria, in contrast, proliferate during incubation: even the lowest inoculum (10^5^ AFU/well) may have expand substantially during the experiment, compressing the effective difference between the three nominal dose levels. The stronger dose-response correlations observed for heat-treated preparations are not evidence that heat-treated bacteria are intrinsically more dose-responsive — they are evidence that the input variable is accurately controlled only for heat-treated preparations. Kruskal-Wallis tests confirmed significant overall concentration effects for **11 of 17** cytokines in heat-treated conditions when tested on condition means (n = 10 strains per dose level); cytokines not reaching significance (IFN-γ, IL-7, IL-12p70, IL-13, IL-17A, MCP-1) showed either consistently low absolute responses or high inter-strain variability across dose levels, providing independent validation of the Spearman correlation findings. This finding is discussed further in Section 4.

#### 3.3.1 Strain-Specific Dose-Response Profiles

Strain-specific dose-response heatmaps revealed marked heterogeneity in dose-sensitivity patterns across the 10 strains (Figure 4). Strains A and B showed monotonically increasing responses (ρ = +1.0) for the majority of cytokines in both viability states, indicating robust dose-dependency regardless of preparation type. In contrast, Strain E showed near-absent dose-response in live conditions (mean ρ = 0.12 across 14 cytokines), suggesting that its immunomodulatory effect saturates at lower bacterial concentrations. Heat-treated preparations demonstrated stronger dose-response than viable preparations for the majority of strains, though the magnitude of this difference was strain-dependent, ranging from negligible (Strains A, I, L) to substantial (Strains B, D, E).

**Figure 4.**
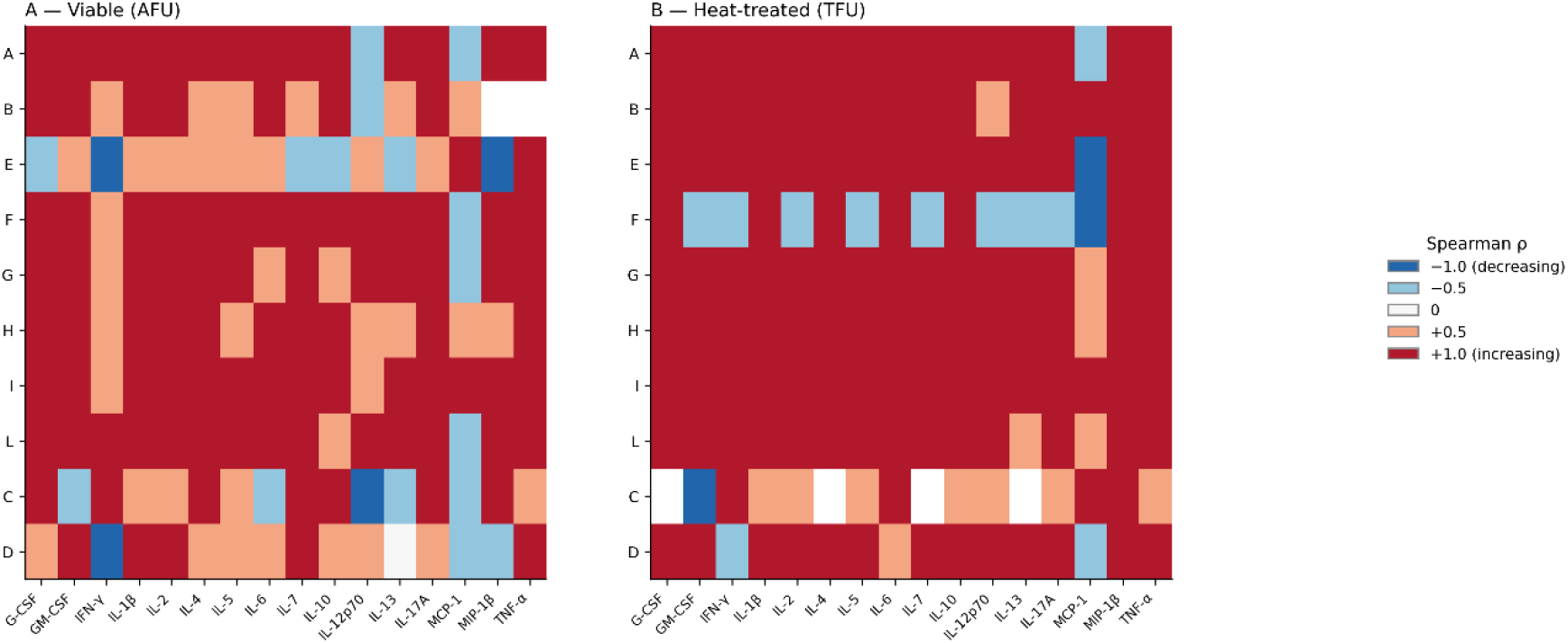
Within-strain dose–response Spearman ρ — categorical representation (IL-8 excluded). Spearman rank correlation between bacterial concentration (10⁵, 10⁶, 10⁷ cells/well) and cytokine fold-change for each strain, calculated on condition means (n = 3 dose-level means per strain per viability state; IL-8 excluded). (A) Viable preparations. (B) Heat-treated preparations. With exactly three ordered dose levels, Spearman ρ is constrained to five discrete values only: −1.0, −0.5, 0.0, +0.5, and +1.0; a categorical colour scale is used accordingly. Dark green (ρ = +1.0): monotonically increasing dose-response across all three concentrations. Dark red (ρ = −1.0): monotonically decreasing. Grey (N/A): all values at LLOQ floor (Strain C heat-treated). Across the panel, 58% of strain–cytokine combinations showed monotonically increasing responses in viable preparations versus 73% in heat-treated preparations.

### 3.4 Dose-Response Profiles of Viable Preparations

Per-strain dose-response profiles of viable preparations are summarized in Figure 5 and reported in full in the Supplementary Material (Section S3).

**Figure 5.**
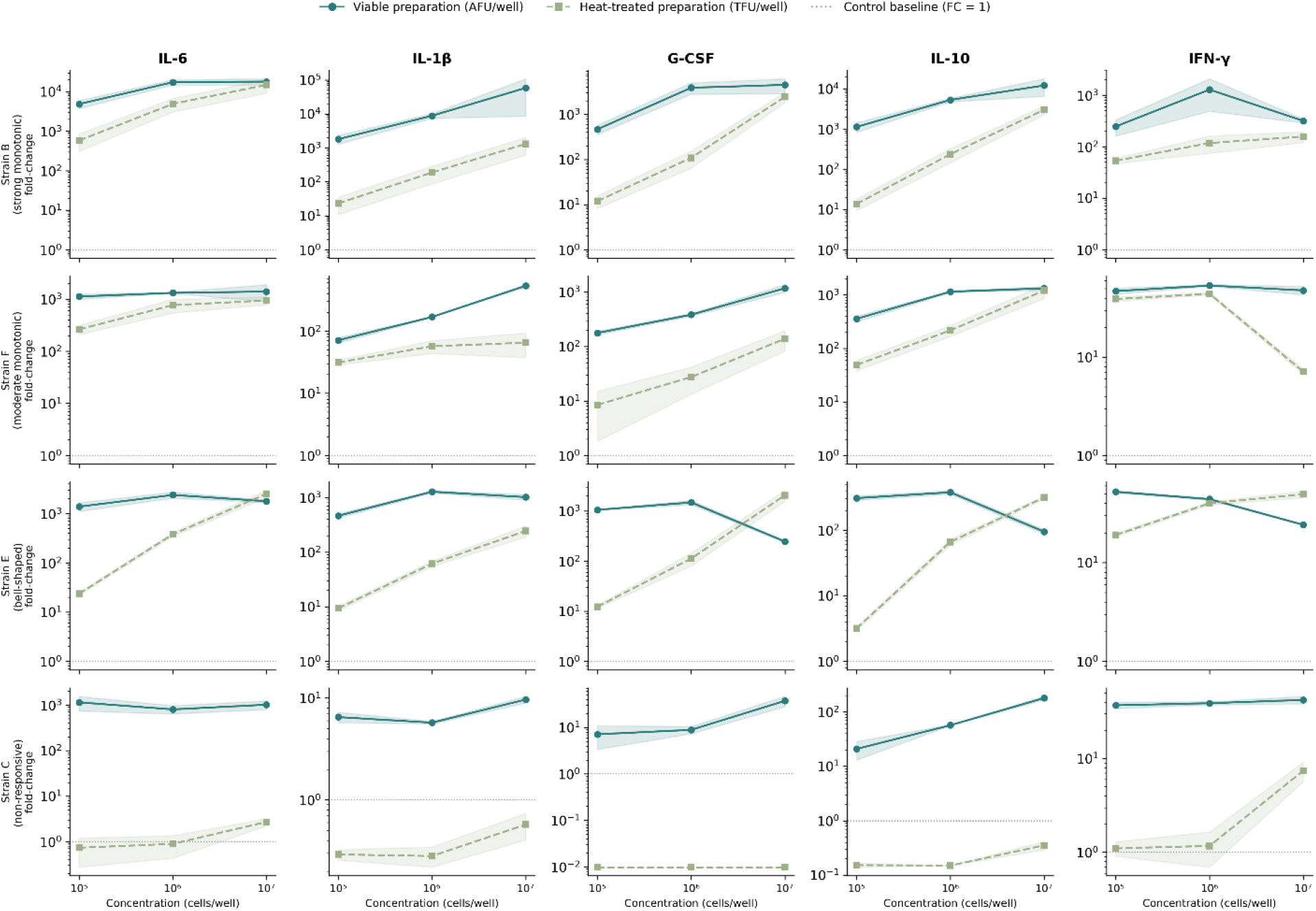
Dose-response profiles of viable and heat-treated preparations across representative strains. Mean fold-change ± SD (n = 3 technical replicates) for five cytokines (IL-6, IL-1β, G-CSF, IL-10, IFN-γ) across 10⁵, 10⁶, 10⁷ cells/well. Solid lines with circles: viable preparations (AFU/well). Dashed lines with squares: heat-treated preparations (TFU/well). Y-axis: log scale. Rows: Strain B (strong monotonic), Strain F (moderate monotonic), Strain E (bell-shaped), Strain C (viability-dependent). Dotted horizontal line: control baseline (fold-change = 1). Values represent means of n = 3 technical replicates; shaded bands represent ±1 SD.

### 3.5 Non-Monotonic Dose-Response Profiles: Bell-Shaped Kinetics

Strains exhibiting non-monotonic, bell-shaped dose-response kinetics are illustrated in Figure 5 and described in detail in the Supplementary Material (Section S4).

### 3.6 Strain C: A Strongly Viability-Dependent Immunomodulatory Profile

The strongly viability-dependent immunomodulatory profile of Strain C, active in the viable state, with attenuated activity after heat treatment, is detailed in the Supplementary Material (Section S5); this observation is examined further in the Discussion as the most consequential finding for postbiotic formulation strategy.

### 3.7 Viable vs Heat-treated: Dose-Dependent Convergence and FED⁷⁰

Systematic comparison of viable (AFU) and heat-treated (TFU) preparations across matched concentration points revealed concentration-dependent convergence of functional activity. At 10⁵ cells/well, HT/V activity ratios for IL-6 ranged from very low (Strain C) to 20% (Strain F). At 10^6^, Strain F HT reached 58% of viable activity. At 10^7^: Strain B HT reached 86% of viable IL-6 (14,980 vs 17,455-fold at the 10⁶ AFU/well reference); Strain E HT exceeded viable at 10^6^ reference (2,512 vs 2,383-fold, 105%); Strain H HT reached 103% at matched dose; Strain A HT at 10^7^ TFU produced 266% of viable IL-6 at the 10^6^ AFU reference.

To operationalize this relationship, we introduce the *Functional Equivalence Dose* (FED^70^) — the minimum TFU/well concentration at which a heat-treated preparation achieves ≥70% of the immunological activity of the corresponding viable preparation at the reference dose of 10^6^ AFU/well. The 70% threshold was selected on two grounds: biologically, it represents a conservative threshold acknowledging that PBMC assay variability — both technical and inter-donor — typically ranges from 20–35% (Li et al., 2016; van Hemert et al., 2010), making this threshold a pragmatic operational criterion for functional equivalence screening in a single-donor context; regulatorily, it represents a conservative adaptation of the 80–125% bioequivalence acceptance range established by EMEA (2010). The FED^70^ matrix (Figure 6) provides a functional basis for postbiotic dose selection currently absent from any regulatory framework. It is noted that for strains exhibiting bell-shaped kinetics (Strains D and E), 10⁶ AFU/well coincides with the peak of the viable dose-response profile. The FED⁷⁰ calculated for these strains therefore represents equivalence to peak viable activity — a more stringent criterion than for monotonically increasing strains, where 10⁶ AFU/well represents an intermediate rather than maximal dose. This asymmetry should be considered when comparing FED⁷⁰ values across strain phenotype groups. Specifically, FED⁷⁰ values for Strains D and E represent equivalence relative to peak viable activity, whereas FED⁷⁰ values for monotonically increasing strains (A, B, F, H, I, L) represent equivalence relative to an intermediate viable dose. These values are therefore not directly comparable across phenotype groups and should be interpreted within each strain’s dose-response context individually.

**Figure 6.**
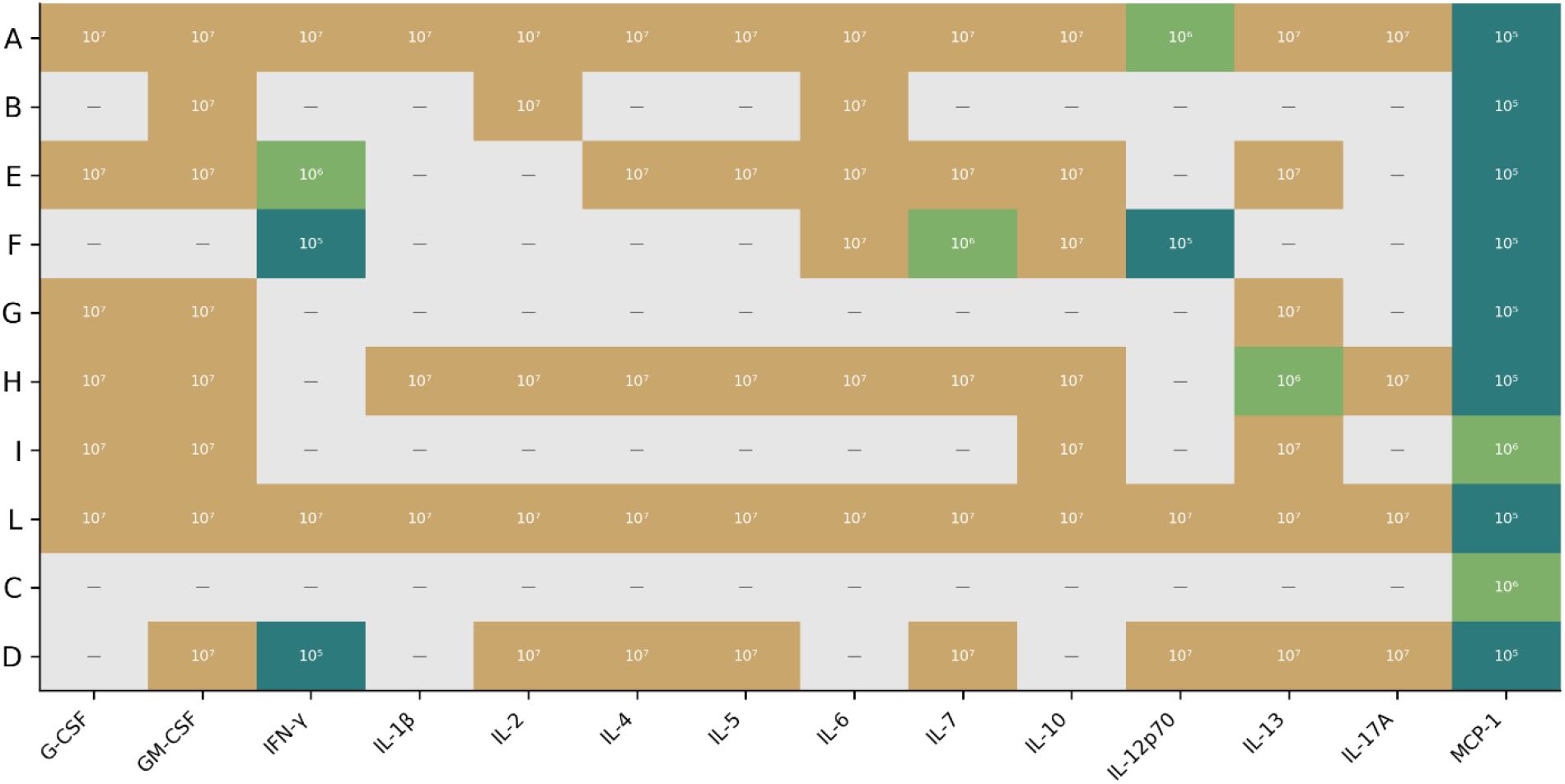
Functional Equivalence Dose (FED⁷⁰) matrix. Minimum TFU/well concentration at which heat-treated preparations achieve ≥70% of the immunological activity of the corresponding viable preparation at the reference dose of 10⁶ AFU/well, for each strain (rows A–L) and 14 quantifiable cytokines (columns; IL-8, MIP-1β, TNF-α excluded). Colour coding: grey = N/A (viable response <1.5× control, equivalence undefined); red = “Not reached” (viable response sufficient but HT did not achieve ≥70% equivalence at any of the three tested doses); light green = FED⁷⁰ at 10⁵ TFU/well; teal = FED⁷⁰ at 10⁶ TFU/well; dark blue = FED⁷⁰ at 10⁷ TFU/well. Data represent means of n = 3 technical replicates.

### 3.8 Cytokine Co-Regulation Networks Reveal Coordinated Response Programs

Inter-cytokine Spearman correlation analysis across the full dataset revealed highly structured co-regulation patterns reflecting coordinated immune response programs. Among the strongest correlations were the Th2/adaptive pairs IL-2 and IL-5 (ρ = 0.97), IL-4 and IL-5 (ρ = 0.95), and IL-2 and IL-4 (ρ = 0.94), reflecting coordinated Th2-type and adaptive immune activation across the dataset. These near-unity correlations across the entire dataset reflect global response amplitude as the dominant source of co-variation: conditions of high bacterial stimulation (viable preparations at high dose) produce uniformly elevated responses across the cytokine panel, while low-stimulation conditions produce uniformly reduced responses. This dataset-level correlation is therefore an ecological phenomenon driven by the viability and dose factors rather than evidence of direct transcriptional co-regulation between individual cytokine pairs. Within-condition correlations, stratified by viability state and dose level, are expected to be substantially lower and would more faithfully represent pathway-specific co-regulation.

The correlation matrix resolved two major clusters. The first comprised TNF-α, IL-1β, IL-6, IFN-γ, IL-2, and IL-12p70, representing coordinated pro-inflammatory and Th1-type responses driven by monocyte/macrophage and T cell activation. The second cluster included IL-4, IL-5, IL-10, and IL-13, reflecting Th2-type adaptive immune responses. G-CSF and GM-CSF formed a partially independent sub-cluster associated with myeloid cell proliferation and differentiation. IL-8 and MIP-1β correlated strongly with each other and with the pro-inflammatory cluster. MCP-1 showed the most independent expression pattern, with substantially lower correlations to both major clusters (mean ρ with pro-inflammatory cluster = 0.25; mean ρ with Th2 cluster = 0.16), consistent with its unique viability-independence demonstrated in Section 3.2 and its operation through distinct regulatory pathways responding to bacterial structural presence rather than metabolic activity. Analysis of cytokine co-regulation profiles across doses revealed that increasing concentration alters the qualitative composition of the immunological signature in a strain-specific manner (Figure 7). For Strain B, non-isometric expansion was observed: IL-6, IL-1β, IL-10, and G-CSF scaled dramatically while IFN-γ, IL-7, and IL-17 showed modest dependence. Strain E showed qualitative remodelling at 10^7^ AFU/well: G-CSF collapsed in the viable form (1,480 → 246-fold) while the heat-treated form produced 2,059-fold — an analyte dominance inversion visible only through multiplex profiling across both forms and the full concentration range. It is noted that the correlation matrix reflects the full dataset spanning all viability states and dose levels, and therefore captures global response amplitude as a major co-variation source. Cytokine pairs that appear strongly correlated at the dataset level may not be co-regulated within individual conditions — a distinction relevant for mechanistic interpretation.

**Figure 7.**
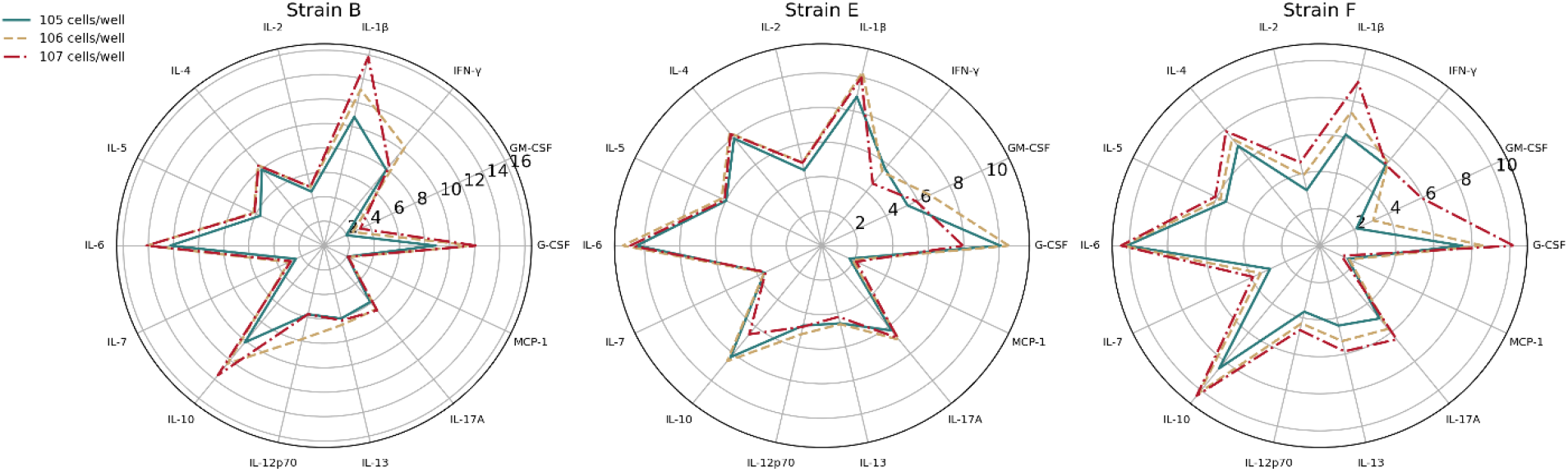
Dose-dependent cytokine co-regulation profiles (viable preparations). Radar charts for Strains B, E, and F across viable dose levels (10⁵, 10⁶, 10⁷ AFU/well). Spokes represent individual cytokines (14 quantifiable analytes); radial distance represents log₂ fold-change versus unstimulated control, clipped at zero (negative values not shown). Concentric rings correspond to log₂ fold-change values of 3, 6, and 9 (i.e. linear fold-changes of 8×, 64×, and 512× versus unstimulated control). Polygons: light shading = 10⁵ AFU/well; medium shading = 10⁶ AFU/well; dark teal = 10⁷ AFU/well. Values represent means of n = 3 technical replicates. Strain B illustrates non-isometric dose-dependent expansion with IL-1β and IL-10 scaling more steeply than IFN-γ; Strain E illustrates bell-shaped kinetics with collapse of G-CSF at 10⁷; Strain F illustrates moderate monotonic escalation.

### 3.9 Strain-Specific Immunomodulatory Signatures Emerge From Data-Driven Analysis

To characterize the immunological fingerprint space at the strain level, PCA was performed on the matrix of condition means (60 conditions: 10 strains × 2 preparation types × 3 doses), collapsing technical replicates to their mean — a distinct analysis from the full-dataset PCA reported in Section 3.10 which preserves replicate-level variance structure.

PCA of the condition-level matrix (60 conditions: 10 strains × 2 preparation types × 3 doses) revealed that the first two principal components explained 85.2% of total variance (PC1: 75.5%, PC2: 9.7%), demonstrating that the 14-cytokine immunological space is highly structured (Figure 8A). Unsupervised hierarchical clustering of strain-level mean fingerprints (Ward linkage, Euclidean distance) identified three functionally distinct clusters (Figure 8B): Cluster 1 — Strains A, B: high-magnitude, broad-spectrum pro-inflammatory cytokine induction; Cluster 2 — Strains E, F, G, H, I, L: moderate responses with diverse kinetic profiles; Cluster 3 — Strains C, D: low-magnitude viable responses and strongly attenuated immunomodulatory activity in the heat-treated form. These cluster assignments reflect the immunological profile of a single donor and should be interpreted as hypothesis-generating phenotypic groupings requiring validation across independent donors.

**Figure 8.**
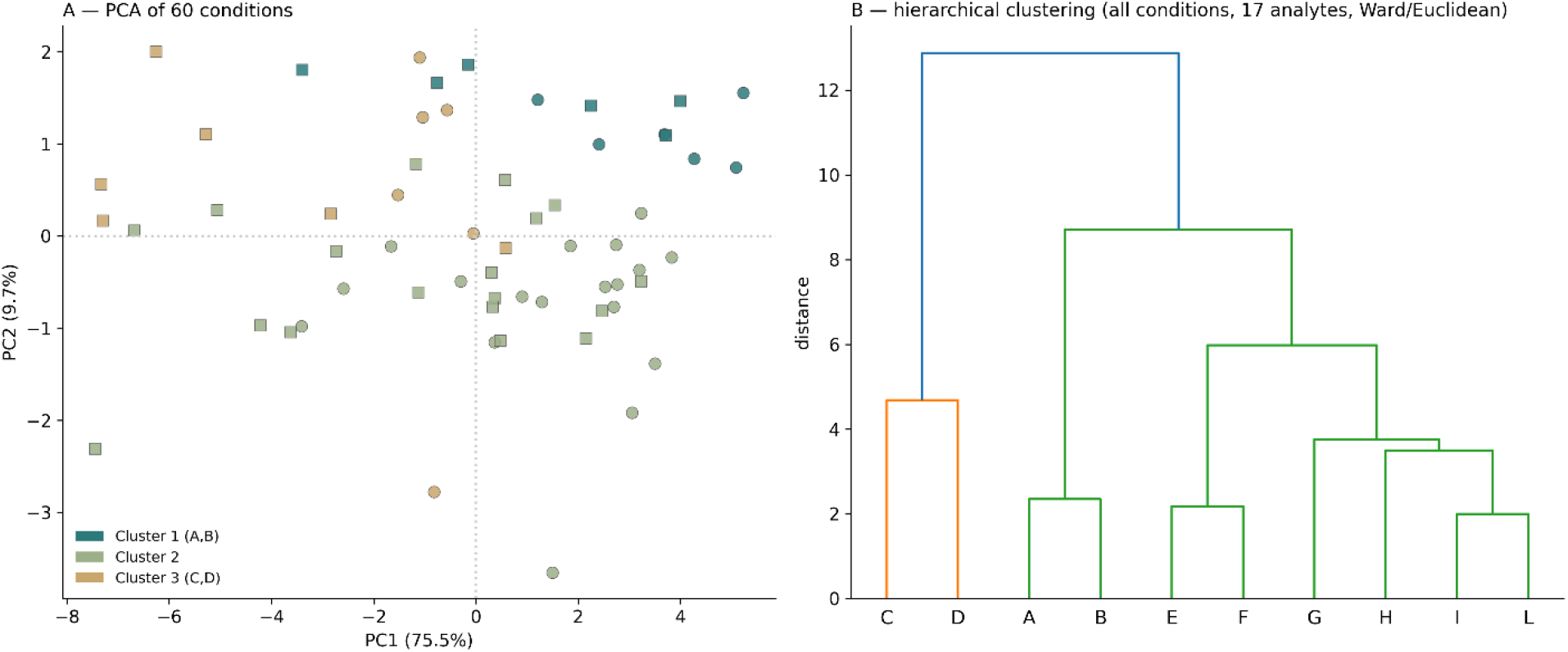
PCA and hierarchical clustering of probiotic immunological fingerprints. (A) PCA of all 60 conditions (10 strains × 2 preparation types × 3 doses). PC1: 75.5% variance; PC2: 9.7% (total 85.2%). Circles: viable (AFU); squares: heat-treated (TFU). Marker size proportional to dose. Strain labels within markers; colours by strain. (B) Hierarchical dendrogram (Ward linkage, Euclidean distance) of strain-level fingerprints across all conditions (17 analytes). Three clusters: Cluster 1 (high-magnitude: A, B); Cluster 2 (moderate: E, F, G, H, I, L); Cluster 3 (low-magnitude: C, D). Leaf label colours correspond to cluster assignment.

Z-score heatmap and functional radar plot analysis revealed distinct immunomodulatory profiles that were not predefined but emerged entirely from the data. In this single-donor model, Strain B exhibited the most potent immunostimulatory profile: TNF-α z-score = +0.51, IL-1β = +1.88, IL-6 = +1.90, IFN-γ = +1.68, with the strongest Th1 bias among high-magnitude strains (Th1/Th2 functional score ratio = 5.1). Strain A showed a similar but less pronounced pro-inflammatory profile (TNF-α z = +0.51, IFN-γ = +1.12, IL-6 z = +1.62, Th1/Th2 = 3.0).

Strains E, F, G, H, I, and L clustered together with moderate pro-inflammatory responses and diverse Th1/Th2 polarization (ratios: 0.5–1.3), with Strains G, H, and I showing Th2-biased profiles (Th1/Th2 = 0.5–0.8) and Strains E and F showing near-balanced activation (Th1/Th2 = 1.0–1.3). Strains C and D exhibited low-magnitude profiles; Strain D showed a balanced Th1/Th2 ratio (1.8) while Strain C showed a markedly viability-dependent profile, with substantially attenuated immunomodulatory activity in the heat-treated form, consistent with its unique thermolabile phenotype. The absolute z-score values and Th1/Th2 ratios reported reflect this donor’s immune background and are expected to vary in absolute magnitude across donors, though relative rankings are anticipated to be preserved.

The inter-strain variation for individual cytokines was substantial: IL-12p70 varied more than 40-fold between the highest-producing strain (B) and the lowest (D) at 10⁷ AFU/well (viable preparations), consistent with the 16-fold variation reported by van Hemert et al. (2010) for L. plantarum strains. Functional cluster classification (3-class, chance 33.3%) achieved 83.3% LOO partition-recovery consistency — 2.50× above chance. Because the three clusters were defined by unsupervised hierarchical clustering of the same cytokine data used to train and evaluate this classifier, this metric reflects internal consistency only and should not be interpreted as expected accuracy for classifying novel strains into these phenotype groups. Binary preparation-type classification (viable vs heat-treated, chance 50%) achieved 63.3% LOO accuracy, confirming that a preparation’s immunological identity is determined jointly by strain origin and viability state.

Random Forest classification of 10 strains from viable preparations (30 conditions, LOO-CV) achieved 40.0% accuracy versus 10% chance --- a4.00× improvement (Figure9A). Feature importance analysis (Figure 9B) identified IL-12p70 (importance = 0.110), IL-13 (0.103), MCP-1 (0.090), IFN-γ (0.090), and IL-7 (0.079) as the cytokines carrying the greatest discriminative information for strain classification. Notably, IL-13 ranked as the second strain discriminator despite not showing a statistically significant difference between viable and heat-treated preparations at the condition mean level (Mann-Whitney FDR p = 0.84; Section 3.2). This apparent contradiction reflects the distinction between two different classification questions: IL-13 does not distinguish viability states globally, but carries strain-specific information — levels characteristic of each ceppo are preserved regardless of preparation type. An analyte can be viability-insensitive yet strain-informative, and this dissociation is precisely what multiplex ML analysis is designed to detect.This ranking diverges from the analytes with the highest absolute fold-changes (IL-6, IL-1β, G-CSF) and is consistent with the IL-10/IL-12 polarization framework validated by Foligné et al. (2007) and O’Mahony et al. (2005): the cytokines that distinguish strains are those reflecting immune polarization direction, not overall activation magnitude.

**Figure 9.**
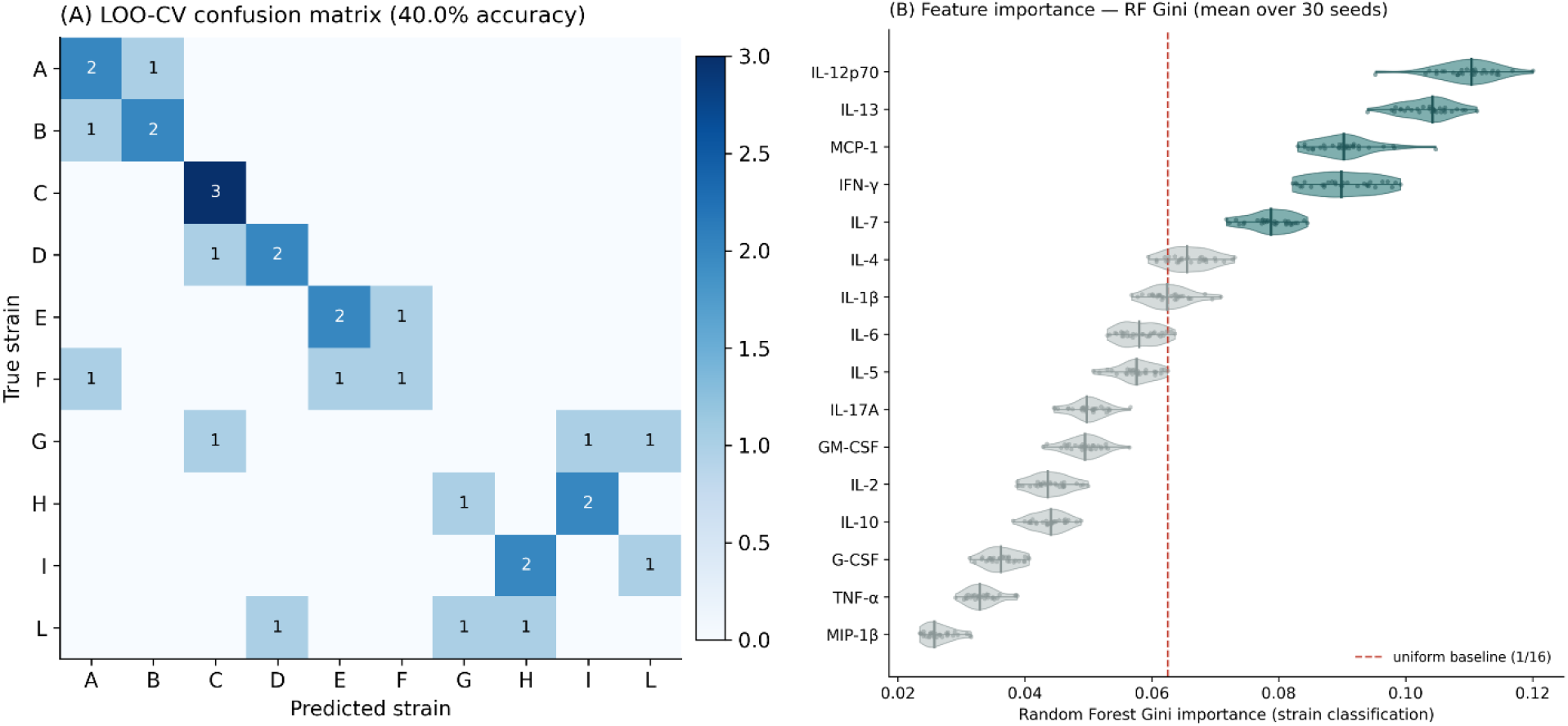
Random Forest strain classification and feature importance (IL-8 excluded). (A) LOO-CV confusion matrix for 10-strain classification of viable preparations (30 conditions; chance level 10%). Classification performed on 16-cytokine profiles (IL-8 excluded due to upper-quantification-limit saturation in the majority of viable conditions). Overall LOO-CV accuracy: 40.0% (4.0× above chance). Rows: true strain; columns: predicted strain. (B) Cytokine feature importance (mean decrease in impurity ± SD across 30 independent model fits). Top discriminators: IL-12p70 (0.110), IL-13 (0.103), MCP-1 (0.090), IFN-γ (0.090), IL-7 (0.079). Dashed red line: uniform baseline (1/16). Top discriminators differ from the highest absolute fold-change cytokines (IL-6, IL-1β, G-CSF), indicating that strain identity information resides in immune polarisation direction, not activation magnitude.

Functional cluster classification (3-class, chance 33.3%) achieved 83.3% LOO accuracy — 2.50× above chance— demonstrating robust separability of the three immunological phenotype groups (Figure 10B). Binary preparation-type classification (viable vs heat-treated, chance 50%) achieved 63.3% LOO accuracy, confirming that thermal treatment produces a consistent, detectable shift in cytokine-induction signature across the strain panel — implying that a preparation’s immunological identity is determined jointly by its strain origin and its viability state. It should be noted that these three clusters were derived by unsupervised hierarchical clustering of the same cytokine data used to train and validate this classifier. The 83.3% accuracy therefore represents an internal consistency metric — confirming that Random Forest can recover the hierarchical partition — rather than an independent validation of the clusters as biological phenotypes. External validation on strains not used to derive the cluster definitions would be required to establish generalisability.

**Figure 10.**
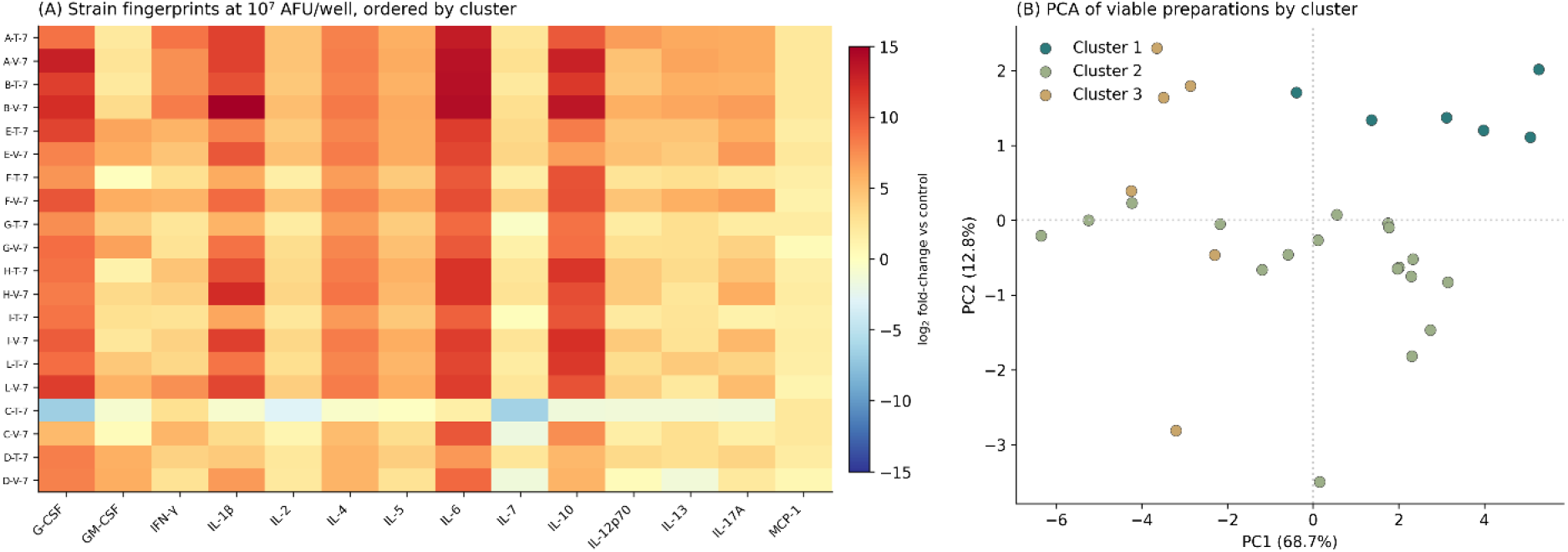
Immunological fingerprint heatmap and cluster validation. (A) Heatmap of strain immunological fingerprints at 10⁷ cells/well, ordered by cluster. Rows: Viable (teal) and Heat-treated (sage) for each strain. Columns: 14 cytokines. Colour: log₂ fold-change (RdYlBu). Heavy lines: cluster boundaries. Cluster 1 (high-magnitude: A, B); Cluster 2 (moderate: E, F, G, H, I, L); Cluster 3 (low-magnitude: C, D). (B) PCA of viable preparations coloured by cluster (Cluster 1: teal; Cluster 2: sage; Cluster 3: red). Marker size proportional to dose. Cluster separation validates biological coherence of the three functional phenotype groups. Overall accuracy: 83.3% (2.50× above chance)

**Figure 11.**
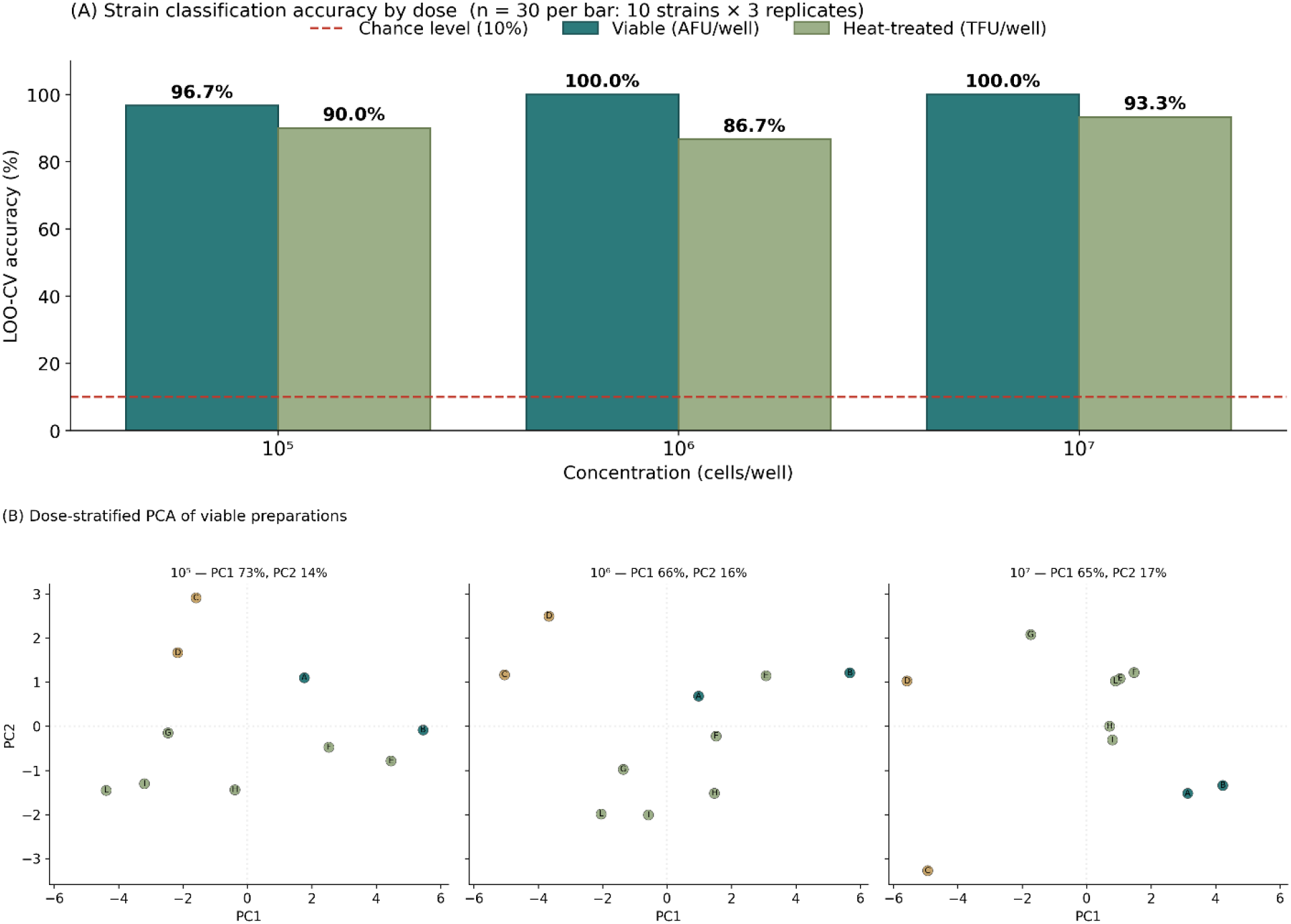
Strain classification accuracy by dose and preparation type (IL-8 excluded). (A) Random Forest LOO-CV accuracy (%) for 10-strain classification at each individual dose level (10⁵, 10⁶, 10⁷ cells/well) for viable (teal) and heat-treated (sage) preparations. Classification performed on 16 cytokines (IL-8 excluded). n = 30 replicate-level samples per bar (10 strains × 3 technical replicates). Dashed red line: chance level (10%). Numbers above bars indicate accuracy percentage. (B) Dose-stratified Principal Component Analysis of viable preparations, illustrating the geometric basis of the dose-dependent classification accuracy shown in (A). Three independent PCAs were performed on the 10 viable strain fingerprints at each dose level (10⁵, 10⁶, 10⁷ cells/well), with each panel reporting its own PC1/PC2 explained variance. Markers are coloured by functional cluster: teal = Cluster 1 (A, B); sage = Cluster 2 (E, F, G, H, I, L); red = Cluster 3 (C, D). Progressive inter-cluster separation across increasing doses provides the immunological signal that drives the increase in classification accuracy from 96.7% at 10⁵ to 100% at 10⁶ and 10⁷.

### 3.10 A Single Principal Component Captures the Dominant Variance Axis

To characterize the global variance structure of the complete dataset including technical replicates, PCA was performed on all 192 observations — complementing the condition-mean PCA reported in Section 3.9 which addressed strain-level fingerprint organization. Principal component analysis on the full dataset (180 observations × 17 cytokines, RobustScaler-standardized log-transformed data) revealed a highly concentrated variance structure. The first principal component explained 74.7% of total variance, implying that a single dominant biological process — overall immune activation magnitude — drives the majority of observed variation. PC1 was loaded by multiple pro-inflammatory and adaptive cytokines, with IL-6 (loading = 0.383), IL-4 (0.322), TNF-α (0.289), IL-2 (0.286), and IL-12p70 (0.283) as the strongest contributors, reflecting the coordinated nature of the cytokine response rather than dominance of a single analyte. PC2 (8.8% variance) was dominated by MCP-1 (loading = 0.774), representing the chemokine axis orthogonal to overall immune activation captured by PC1. PC3 (5.3% variance) captured additional myeloid activation variance associated with GM-CSF and G-CSF. Cumulatively, three components explained 89.5% and five components explained 92.7% of total variance, demonstrating that the 17-dimensional cytokine space can be effectively represented by a low-dimensional manifold. Score plots coloured by viability status demonstrated clear separation between viable and heat-treated samples along the PC1 axis: viable bacteria samples occupied the positive PC1 space (higher immune activation), while heat-treated samples were displaced toward the negative end. Within each viability group, concentration-dependent gradients were visible, with 10^7^ AFU/well conditions positioned at higher PC1 values than 10^5^ conditions. This visual separation foreshadowed the high classification accuracy achieved by the SVM algorithm (Section 3.12).

### 3.11 Unsupervised Clustering Validates Viability-Dependent Response Patterns

Both silhouette score and Calinski-Harabasz index optimization identified **k = 2** as the optimal cluster number across k = 2 to 10, strongly suggesting that the dominant partition in the data corresponds to a binary variable — consistent with the viability state factor. K-means clustering at k = 2 revealed a strong but imperfect association between cluster membership and bacterial viability status. Cluster 1 (n = 54) comprised predominantly heat-treated samples (77.8% heat-treated, 22.2% viable), characterized by lower overall cytokine activation magnitude. Cluster 2 (n = 126) was enriched for viable bacteria samples (61.9% viable, 38.1% heat-treated) with higher pro-inflammatory cytokine expression, particularly IL-6, IL-8, MIP-1β, and TNF-α. The imperfect alignment between clusters and viability states is biologically informative. The viable samples assigned to the predominantly heat-treated Cluster 1 were exclusively from the lowest concentration (10⁵ AFU/well), where immune response magnitude approaches that of heat-treated preparations. Conversely, heat-treated samples in Cluster 2 were distributed across 10⁵ (n=5), 10⁶ (n=17), and 10⁷ (n=26) TFU/well, reflecting that a subset of heat-treated preparations at mid-to-high concentrations achieves sufficient PRR engagement to cluster with viable samples. This overlap region identifies the concentration range where immunological distinction between viable and heat-treated preparations becomes ambiguous, providing empirical boundaries for the dose-dependent transition between viability-dependent and viability-independent immunomodulation. Hierarchical clustering using Ward’s method corroborated the k = 2 partition and additionally revealed sub-structure within each cluster corresponding to concentration levels, with the cluster centroid profiles confirming that the two clusters differ primarily in magnitude of pro-inflammatory cytokine production rather than qualitative cytokine pattern.

### 3.12 Support Vector Machine Classification Achieves Near-Perfect Discrimination

Four classification algorithms were evaluated for binary discrimination of viable from heat-treated bacterial responses based on 17-cytokine profiles, with hyperparameters optimized by grid search and 5-fold stratified cross-validation. When classification was performed on condition means (n = 60, LOO-CV) with IL-8 excluded from the feature set due to upper-quantification-limit saturation in the majority of viable conditions (see Section 3.1), Random Forest achieved AUC-ROC 0.702, accuracy 63.3%. This performance, while above chance (50%), is substantially lower than estimates derived from replicate-level analysis, reflecting both the reduction in effective sample size when using condition means and the removal of the ULOQ-saturation signal carried by IL-8. XGBoost achieved AUC-ROC 0.664 (accuracy 65.0%). Support Vector Machine with radial basis function kernel achieved AUC-ROC 0.517 (accuracy 50.0%). Logistic Regression achieved AUC-ROC 0.674 (accuracy 61.7%); the Support Vector Machine was thus the weakest classifier, performing at chance level (Supplementary Figure 3). The superiority of the tree-based ensemble indicates that the decision boundary separating viability states in cytokine space benefits from the conditional splitting used by tree-based methods. McNemar’s test confirmed statistically significant superiority of the Random Forest over SVM (p = 0.004) and Logistic Regression (p < 0.001); the difference between Random Forest and XGBoost did not reach statistical significance (p = 0.080). Feature importance analysis revealed consistent cross-algorithmic identification of key discriminative cytokines. IL-6 emerged as the top discriminative feature across classifiers: SVM permutation importance 0.037, RF Gini importance 0.143, consistent with its dominant loading on PC1 and its role as an integrator of PRR signaling. In Random Forest: IL-6 (0.143), IL-1β (0.081), MIP-1β (0.078). In Logistic Regression: IL-6, MIP-1β, IL-4.The convergence of feature importance rankings across kernel-based (SVM), tree-based (XGBoost, RF), and linear (LR) approaches provides robust evidence that the identified cytokines represent genuine biological discriminators. Misclassified samples (∼32% of total) were concentrated at the dose extremes: 10⁵ AFU/well viable samples misclassified as heat-treated (insufficient viable-metabolism signal at low dose) and 10⁷ AFU/well heat-treated samples misclassified as viable (sufficient MAMP mass at high dose mimicking viable magnitude). This systematic misclassification pattern reinforces the biological interpretation of the clustering overlap described in Section 3.11.

## 4. Discussion

This study was designed to address four interconnected methodological questions that, despite their foundational importance, remain inadequately answered in the probiotic literature: (1) are there measurable and reproducible differences in immunomodulation between live and heat-treated bacteria? (2) can dose-response relationships be demonstrated across a range of concentrations? (3) how does the method of bacterial quantification affect the reliability of dose-response data? and (4) can machine learning tools extract biologically meaningful information from a well-structured factorial experimental design? The results provide clear affirmative answers to all four questions, while simultaneously exposing the methodological fragility of conventional approaches in the field and raising a fifth question — why the PBMC co-culture model, despite its known limitations, remains the most translationally valid platform for probiotic immunomodulatory screening.

### 4.1 The Question of Viability: Live Versus Heat-Treated Bacteria

The 356.9-fold difference in TNF-α secretion between live and heat-treated conditions is, to our knowledge, among the largest fold-changes reported for a single cytokine in a systematically controlled PBMC-probiotic co-culture comparison. Rather than simply confirming that live bacteria are “more potent,” this magnitude of difference demands mechanistic interpretation and raises critical questions about experimental design. The 16 of 17 significantly affected cytokines demonstrate that the viability effect is not pathway-specific but represents a global amplification of immune activation. This is consistent with at least four complementary mechanisms unique to living bacteria. First, metabolically active bacteria continuously release MAMPs through cell division, secretion, and autolysis, creating a sustained and amplifying signal rather than the fixed bolus of ligands presented by an heat-treated preparation. Second, live bacteria produce immunologically active metabolites — short-chain fatty acids, lactic acid, hydrogen peroxide, and bacteriocins — that constitute additional stimulatory signals absent from heat-treated preparations (Lebeer et al., 2010). Third, certain PRR pathways may be preferentially activated by viability-dependent structures, such as bacterial messenger RNA recognized by TLR8, which is rapidly degraded in dead cells but continuously synthesized in living bacteria. Fourth, acidification of culture medium by live bacterial metabolism may modulate immune cell function independently of receptor-mediated signaling.

However — and this constitutes the central interpretive hypothesis of our argument, one that is consistent with but not directly proven by the present data — these same mechanisms that make live bacteria more immunostimulatory also make them uncontrollable as an experimental variable. During a 24-hour co-culture at 37°C in nutrient-rich RPMI medium, live bacteria proliferate. The bacteria-to-PBMC ratio at experiment’s end is unknown and unknowable without endpoint enumeration, which was not performed in this study and is rarely performed in any published study. A conservative estimate — assuming one bacterial doubling every 60–90 minutes for the first 6–8 hours before nutrient depletion or immune-mediated killing slows growth — suggests that the initial inoculum at 10^7^ AFU/well could have expanded by 64- to 256-fold before reaching stationary phase. This means that comparing ‘live 10^5^’ with ‘live 10^7^’ is not a comparison across a 100-fold concentration range — it may be a comparison across a much narrower effective range once proliferation equalizes the groups. Direct testing of this hypothesis requires endpoint bacterial enumeration — a measurement not performed in this study and rarely performed in any published probiotic co-culture study.

This interpretation is directly supported by our dose-response data. Heat-treated preparations, whose concentration remains fixed throughout the experiment, demonstrated significantly stronger Spearman dose-response correlations (mean within-strain ρ up to 1.00 for G-CSF, IL-4, and MIP-1β) than viable preparations for 13 of 17 cytokines — despite inducing lower absolute responses. This is consistent with — though not proven by — the hypothesis that the input variable is more accurately controlled for heat-treated than for viable preparations. The weaker dose-response correlation in viable conditions is most parsimoniously explained, we argue, as a potential artefact of uncontrolled bacterial proliferation compressing the effective dose range, rather than evidence of a fundamentally different dose-response biology. Alternative explanations — including greater intrinsic biological variability in the live-bacteria response — cannot be excluded without endpoint enumeration data. This finding has a direct implication for the interpretation of all published probiotic PBMC co-culture studies that compare viable bacteria across dose levels without endpoint enumeration: their dose-response data are potentially confounded by unquantified within-experiment concentration drift.

The practical implication is methodological and productive: heat-treated preparations provide a more experimentally tractable model for dose-response characterization precisely because their concentration is stable throughout the experiment. Salminen et al. (2021)’s postbiotic definition thus gains an unexpected dimension: postbiotics are not merely an alternative biological concept but a superior experimental tool for controlled pharmacological investigation. This finding aligns with and extends the clinical postbiotic literature — the landmark Depommier et al. (2019) trial showed pasteurized *Akkermansia muciniphila* outperforming its live counterpart on primary metabolic endpoints, Zorzela et al. (2017) found equivalence in 86% of preventive trials, and Andresen et al. (2020) demonstrated clinical efficacy of heat-treated *B. bifidum* MIMBb75 in irritable bowel syndrome — by adding a mechanistic layer: the apparent immunological superiority of live preparations may be partly illusory, reflecting experimental artefacts from bacterial proliferation rather than genuine pharmacological differences.

#### 4.1.1 Bayesian and Pathway-Level Confirmation

The Bayesian framework adds probabilistic quantification to this viability comparison. While live conditions achieved P(immunostimulatory) ranging from 0.64 to 0.97, heat-treated conditions at 10⁵ TFU/well for Strains C and I yielded P values as low as 0.19, identifying conditions where immunostimulation is genuinely uncertain rather than merely reduced in magnitude. This probabilistic view complements the frequentist fold-change analysis and provides actionable dose thresholds. Pathway mapping further demonstrates that the viability effect is not pathway-selective: all 12 signaling cascades show significantly higher activation in live conditions, confirming a generalized amplification mechanism consistent with continuous MAMP release and active metabolite production by viable bacteria.

### 4.2 The Question of Dose: Systematic Dose-Response Characterization

The answer is unambiguously yes, but with critical caveats tied to both quantification methodology and dose-response shape. Across the three tested concentrations (10^5^, 10^6^, 10^7^ AFU/mL, corresponding to bacteria-to-PBMC ratios of approximately 0.5:1, 5:1, and 50:1), dose-dependent cytokine induction was demonstrated for the majority of the panel in six of ten strains, extending the limited prior dose-response evidence base (Ghadimi et al., 2008; Evrard et al., 2011; Hua et al., 2010) to the first systematic multi-strain framework. This is itself a novel contribution: Ouwehand (2017) concluded that the probiotic field lacks dose-response evidence for immune endpoints; Shane et al. (2010) criticized the prevalence of single-dose designs; and Christensen et al. (2006), in one of the few multi-dose human probiotic immune studies, found no significant immune changes at any of five dose levels in 71 participants — a null result that our data suggest may reflect CFU-based dose uncertainty rather than genuine dose-independence.

The detection of immunologically meaningful responses even at the lowest concentration (10^5^ AFU/mL) challenges the widespread assumption that a minimum of approximately 10^6^ bacteria per milliliter is required for immune effects. While responses at 10^5^ AFU/mL were quantitatively lower than at higher concentrations, they were consistently above negative control levels for most cytokines in both viability states. This observation has practical significance for the postbiotic field, where delivered doses may be lower than those achievable with viable preparations due to formulation constraints.

However, the assumption of monotonic dose-response scaling cannot be generalized across all strains. The non-monotonic profiles of Strains D and E — with 83% G-CSF and 25% IL-6 reduction at 10^7^ versus 10^6^ AFU/well — are consistent with the hormesis concept proposed by Hua et al. (2010) and the dosage optimum identified by Ghadimi et al. (2008). Several non-mutually exclusive mechanisms could account for this pattern. The most parsimonious candidate involves IL-10-mediated autocrine feedback: substantial IL-10 induction at 10⁶ could create a suppressive milieu reducing IL-6 and G-CSF output at 10⁷ through STAT3/SOCS3-mediated NF-κB suppression (Latvala et al., 2011). TLR desensitization upon sustained high-density PAMP exposure and induction of negative innate signaling regulators (IRAK-M, A20/TNFAIP3, TOLLIP) represent alternative or complementary candidates (Akira and Takeda, 2004). The present single-timepoint design does not permit mechanistic discrimination between these hypotheses — time-course experiments with IL-10 neutralization or TLR pathway inhibitors would be required to establish causality. The dissociation between viable and heat-treated attenuation patterns at 10^7^ argues against non-specific cytotoxicity and in favour of an active-cell-dependent mechanism. The translational consequence is direct: for these strains, dose escalation beyond 10^6^ AFU/well does not improve immunological efficacy and reduces it for multiple analytes — directly challenging the industry practice of dose maximization that Shane et al. (2010) and Ouwehand (2017) have criticized on evidentiary grounds.

From a regulatory perspective, these data demonstrate that the in vitro PBMC co-culture system, when combined with appropriate bacterial enumeration and statistical analysis, can generate the type of dose-response evidence that regulatory bodies require but the field has largely failed to produce. The EFSA requirement that clinical evidence must be conducted “at the same dosage” as the marketed product, and the ICH E4 mandate for at least three dosage levels in dose-finding studies, reflect pharmacological standards that are achievable in probiotic research but have been systematically neglected. The concentration-matched AFU/TFU comparison enabled by flow cytometric quantification further permits, for the first time, the determination of a Functional Equivalence Dose — the concentration at which heat-treated preparations achieve a defined percentage of the viable preparation’s immunological output. This FED concept, adapted from the biosimilar regulatory framework (EMEA, 2010), provides a strain-specific, endpoint-specific benchmark for postbiotic dose justification that the field currently lacks entirely.

#### 4.2.1 Strain-Specific Dose-Response Heterogeneity

The strain-specific dose-response heatmaps reveal an additional layer of complexity not captured by aggregate analysis. Strains A and B maintain robust dose-dependency in both viability states, while Strain E shows near-absent dose-response in live conditions, with the majority of cytokines showing non-monotonic or flat patterns (ρ = 0 or ρ = −0.5)— suggesting — if the proliferation hypothesis holds — that dose compression may be strain-dependent, potentially reflecting differences in growth kinetics under co-culture conditions. For heat-treated preparations, the systematically stronger and more linear dose-response across all strains provides a practical recommendation: dose-optimization studies should preferentially use heat-treated preparations to obtain interpretable pharmacological data unconfounded by inter-strain variation in growth rate.

### 4.3 The Question of Measurement: Accurate Bacterial Quantification

The choice of bacterial enumeration method is not a technical footnote — it is a determinant of whether dose-response data are interpretable. This study employed flow cytometry enumeration according to ISO 19344, with results expressed in active fluorescent units for viable preparations and total fluorescent units for heat-treated preparations. Three dimensions of this choice merit emphasis. First, flow cytometry provides substantially greater precision than plate counting:

Tracey et al. (2023) quantified the relative standard deviation of flow cytometry at 6.70% versus 12.52% for conventional plate counting (P = 0.0002), with viable cell recovery of 101.8 ± 6.95% by flow versus 81.37 ± 16.03% by plate count. In practical terms, the three concentration levels in our study are known with approximately 7% coefficient of variation; with plate counting, the same nominal concentrations could vary by 15–30%, substantially reducing the statistical power to detect concentration-dependent effects. The Christensen et al. (2006) null result in a five-dose human study may thus reflect dose uncertainty as much as genuine dose-independence.

Second, and critically for the live versus heat-treated comparison at the heart of this study, flow cytometry is the only enumeration method capable of verifying dosing equivalence between the two viability states. Colony-forming unit plate counting yields zero for heat-treated preparations by definition; optical density measurements cannot distinguish intact from lysed cells. Flow cytometry, by detecting individual intact fluorescently labelled cells regardless of metabolic activity, provides direct verification that equivalent numbers of intact bacterial cells are present in both preparations — a standard that CFU-based approaches fundamentally cannot meet for heat-treated material.

Third, the regulatory significance of this methodological choice is growing. The European Food Safety Authority approved pasteurized *A. muciniphila* as a novel food in 2021 based on total fluorescent unit enumeration by flow cytometry rather than colony-forming unit counting (EFSA, 2021), establishing a precedent that may reshape how probiotic and postbiotic doses are reported. We note with concern that the vast majority of published probiotic immunomodulation studies continue to rely on colony-forming unit plate counting or optical density-based estimates. When the input variable of a dose-response experiment is known only within an order of magnitude, the experiment cannot distinguish dose-dependent from dose-independent effects — regardless of statistical sophistication applied to the output.

### 4.4 The Question of Analysis: Machine Learning and Functional Annotation

The fourth pillar of this study addresses a methodological opportunity rather than a limitation: the use of machine learning to extract information from well-structured experimental data that goes beyond the original experimental hypothesis. The ML analysis reported here is, to our knowledge, the first application of supervised and unsupervised classification to probiotic strains based on their immunomodulatory cytokine-induction phenotype. The classification accuracies reported reflect descriptive pattern recognition within a single-donor, technical-replicate dataset and should not be interpreted as inferential estimates of generalization to novel donors or independent experimental runs. Charizani et al. (2024) described the first supervised ML model for probiotic potential prediction from in vitro phenotypic features, but used cell surface properties in an aquatic model. Coelho and Mossotto et al. (2020) applied hierarchical clustering to PBMC NOD2/TLR stimulation cytokine data for patient stratification — the closest methodological precedent in the immunological context, applied to a different classification goal. Wu et al. (2024) differentiated 12 Bifidobacterium strains by ML on intestine-on-a-chip readouts rather than cytokine profiles. The present study bridges these parallel trajectories: genomic ML (Orkkatteri Krishnan et al., 2025; Sun et al., 2022; Shufang et al., 2024), clinical cytokine ML (Wandong et al., 2022; Li P et al., 2025), and functional probiotic phenotyping (van Hemert et al., 2010; Meijerink et al., 2010) — extending the Random Forest gene-trait matching of the latter into the immunological phenotype space itself.

The Random Forest classifier discriminated live from heat-treated responses with AUC-ROC 0.702 (condition-level LOO-CV, IL-8 excluded), establishing that the 17-cytokine profile contains sufficient information to infer bacterial viability status with high accuracy. It is important to distinguish this external-facing metric (AUC-ROC 0.702 for viability classification, trained and tested on non-overlapping partitions) from the 83.3% cluster-classification accuracy reported in Section 3.9, which is an internal consistency metric reflecting partition recovery rather than generalization performance. The convergence of feature importance rankings across algorithmically diverse classifiers — kernel-based SVM, tree-based XGBoost and Random Forest, linear Logistic Regression — provides a form of computational cross-validation that strengthens biological inference. Following exclusion of IL-8 from the feature set due to upper-quantification-limit saturation — which would otherwise act as a binary ULOQ flag rather than a genuine biological discriminator — IL-12p70, IL-13, and IFN-γ emerged as top discriminators for strain classification, rather than IL-6 or IL-1β. This finding is biologically coherent with the Foligné et al. (2007) IL-10/IL-12 validation and the clinical correlation of O’Mahony et al. (2005): the most informative immunological dimension for strain identity is not activation magnitude but polarization direction and pattern specificity. An interpretive consideration warranting explicit attention concerns the 17-plex panel’s inclusion of analytes typically associated with adaptive immune responses — IL-2, IL-4, IL-5, IL-13, IL-17. Their presence in a 24-hour PBMC stimulation assay does not reflect probiotic-driven adaptive immune polarization; rather, it reflects bystander activation of resting memory T cells and NK cells by monocyte-derived IL-12p70 and TNF-α (Akira et al., 2006) and background secretion from basophil precursors. IL-12p70 emerged as the top-ranked discriminator for strain classification (importance = 0.110), with IL-13 second (0.103); MCP-1 ranked third (0.090), consistent with its unique regulatory properties demonstrated in Section 3.2. This demonstrates the value of unbiased multiplex panel selection over hypothesis-driven single-analyte screening — restricting the panel to “expected” innate analytes would have missed top-ranking discriminative features.

More provocatively, the machine learning approach revealed functional phenotypes that were not part of the original experimental question. Three distinct immunomodulatory phenotypes emerged entirely from data-driven analysis: high-magnitude/Th1-biased (Th1/Th2 = 5.1 for Strain B), moderate with variable polarization (Th1/Th2 = 0.5–1.3 for Strains E, F, G, H, I, L), and low-magnitude (Strains C, D). The high cluster classification accuracy (83.3% LOO-CV) reflects the internal separability of these groups within this dataset; because the clusters were defined from the same data on which the classifier was trained, this metric should not be interpreted as the expected accuracy for classifying novel strains into these phenotype groups. The identification of these phenotypes among just 10 strains carries a sobering implication for the field. McFarland et al. (2023) documented that probiotic systematic reviews and meta-analyses continue to group strains of the same species as interchangeable. Our data demonstrates that immunomodulatory phenotypes within a small collection range from strongly pro-inflammatory to anti-inflammatory. Pooling such strains in a meta-analysis is pharmacologically equivalent to combining clinical trials of agonists and antagonists acting on the same target and concluding that the drug class has “no effect.”

### 4.5 Model Selection and Translational Validity

The choice of the PBMC co-culture model as the immunological readout platform for this study was not made by default but reflects a deliberate assessment of translational and analytical requirements that no alternative in vitro system can simultaneously satisfy. PBMCs are primary human cells — not immortalized lines carrying oncogenic signaling aberrations. Jurkat T-cell lymphoma cells harbour PTEN and SHIP1 loss resulting in constitutively hyperactivated PI3K/Akt signaling that profoundly distorts T cell receptor-dependent cytokine production (Abraham and Weiss, 2004). THP-1 monocytic leukemia cells exhibit constitutive NF-κB activation, altered TLR1/2/4/6 expression, and quantitatively different cytokine secretion profiles from primary monocytes (Bosshart and Heinzelmann, 2016; Tedesco et al., 2018). These are not minor technical caveats; they are fundamental distortions of the very signaling pathways under investigation.

More critically, PBMCs constitute a heterogeneous population — monocytes, T lymphocytes, B lymphocytes, natural killer cells, and dendritic cells in physiological proportions (Kleiveland, 2015; Maecker et al., 2012). The 17-plex cytokine fingerprint that forms the basis for machine learning classification in this study emerges from the functional cross-talk among these subpopulations: monocytes and dendritic cells produce TNF-α, IL-6, and IL-12 through TLR2 engagement (Trinchieri, 2003); NK cells produce IFN-γ mediated by DC-derived IL-12 through a positive feedback loop (Vivier et al., 2011; Iwasaki and Medzhitov, 2015); T helper cells generate IL-4 and IFN-γ reflecting Th1/Th2 polarization. No single cell line can generate this dimensionality. The systems immunology literature has established that immune responses form coordinated multi-cytokine modules, and that single-parameter measurements fundamentally cannot capture emergent immune properties (Brodin and Davis, 2017; Davis et al., 2017). Intestinal organoids, while increasingly sophisticated, are epithelial structures devoid of immune cells by design (Sato et al., 2009; Clevers, 2016; Bar-Ephraim et al., 2020). Immune-competent organoid co-cultures remain technically immature, low-throughput, and unstandardized (Dijkstra et al., 2018; Neal et al., 2018). Organ-on-a-chip systems offer the most physiologically complete architecture but are fundamentally incompatible with the high-throughput, multi-strain, multi-concentration screening format required by factorial experimental designs (Ingber, 2022).

The decisive argument for the PBMC model, however, is not its superiority in any single dimension but its unique position as the only in vitro platform with documented clinical predictive validation for probiotic immunomodulation. The Foligné et al. (2007) correlation between PBMC-derived IL-10/IL-12 ratios and in vivo anti-inflammatory capacity in murine colitis models was validated clinically by O’Mahony et al. (2005) in irritable bowel syndrome patients, confirmed at the human mucosal transcriptome level by van Baarlen et al. (2011), and translated into clinical trial strain selection in the PandA allergy prevention trial (Niers et al., 2005; Niers et al., 2009). This constitutes the most complete bench-to-bedside validation chain for any in vitro probiotic screening platform. No equivalent predictive validation exists for any cell line model, organoid system, or organ-on-a-chip platform in the context of probiotic immunomodulation (Papadimitriou et al., 2015; Klaenhammer et al., 2012). The inter-donor variability often cited as a limitation of PBMC assays is, in fact, a feature reflecting real human immune diversity: the Milieu Intérieur consortium demonstrated across 1,000 healthy donors that PBMC cytokine variation is driven by genetics, age, sex, and environmental factors (Piasecka et al., 2018; Patin et al., 2018), while the Human Functional Genomics Project showed that gut microbiome composition directly correlates with PBMC cytokine production capacity (Schirmer et al., 2016) — the very diversity that determines differential clinical responses to probiotics.

Our use of a 17-plex panel represents a direct methodological advance over the original two-cytokine Foligné ratio. While the IL-10/IL-12 ratio remains a useful first-pass metric, it has been shown to oversimplify complex cytokine networks (Latvala et al., 2008), and van Hemert et al. (2010) demonstrated that multiplex profiles discriminate strains indistinguishable by any single cytokine pair. The expanded panel, combined with machine learning, captures Th1/Th2/Th17/regulatory axis complexity and enables granular strain discrimination that aligns with the systems immunology paradigm now standard in vaccine research (Tsang et al., 2014; Pulendran et al., 2010) but still underutilized in probiotic characterization. Dong et al. (2012) recommended PBMCs from at least six to ten donors with individual donor data reporting for robust strain ranking; our single-donor design captures strain-level resolution at the expense of population-level generalizability, a trade-off explicitly addressed in the limitations below.

### 4.6 MCP-1 Independence: An Internal Methodological Control

The viability-independence of MCP-1 secretion (p = 0.61, fold-change 1.0 across all strains and concentrations) serves as an internal methodological control that strengthens the validity of all other comparisons. MCP-1 (CCL2) is produced through both MyD88-dependent and MyD88-independent signaling pathways and can be activated by bacterial structural components — peptidoglycan, lipoteichoic acid — independently of viable bacterial metabolism. Its consistent secretion across both viability states argues that PBMCs were exposed to equivalent structural bacterial material in both conditions, confirming that the experimental system is not confounded by preparation-specific differences in cell lysis, endotoxin contamination, or media components.

From an ML perspective, the viability-independence of MCP-1 is equally significant: despite showing no differential expression between viable and heat-treated conditions globally, MCP-1 ranked as the third most important feature for strain classification (importance = 0.090). This dissociation — between MCP-1’s failure to discriminate viability states and its contribution to strain identity discrimination — illustrates a core principle of ML-informed multiplex analysis: the informativeness of a cytokine depends entirely on the classification question. An analyte uniformly insensitive to one variable may be highly informative for another. This finding could not have been identified through hypothesis-driven, single-analyte screening and demonstrates the value of unbiased multiplex ML feature selection.

### 4.7 Limitations

Several limitations warrant transparent acknowledgment. First and most critically, the single-donor design, while enabling precise strain-specific characterization by eliminating inter-individual variability as a confounding factor, limits generalizability to the broader population. Li et al. (2016) characterized cytokine production variability across 197 donors and identified six genome-wide significant cytokine quantitative trait loci, demonstrating a genetic basis for inter-individual variation. The Milieu Intérieur consortium further established that age, sex, and genetic background are major determinants of cytokine response variation (Piasecka et al., 2018; Patin et al., 2018). Both Foligné et al. (2007) and van Hemert et al. (2010) demonstrated that relative strain rankings are consistent across donors despite absolute value differences, suggesting that the ordinal relationships reported here are likely reproducible, but validation across at minimum five donors of diverse sex, age, and immune backgrounds is necessary before population-level conclusions can be drawn. Specifically, the three functional cluster assignments, the Th1/Th2 polarization ratios, the FED⁷⁰ thresholds, and the ML classification accuracies reported here all reflect a single immune background. The ordinal relationships between strains are anticipated to be preserved; the absolute quantitative values are not. Multi-donor validation is required before any of these parameters can be used for population-level dose characterization or regulatory substantiation. The ML classification accuracies derive from cross-validation applied to technical replicates within a single-donor experiment. While the dataset size (60–180 observations per classifier) is adequate for model fitting, the structural correlation between technical replicates of the same condition introduces dependency not fully addressed by cross-validation. External validation on independent biological replicates from additional donors is required to confirm generalizability.

Second, the uncontrolled proliferation of live bacteria during the 24-hour co-culture represents an inherent limitation of all live bacteria co-culture experiments and should be acknowledged explicitly. Despite flow cytometric enumeration at T=0, the bacteria-to-PBMC ratio at experiment’s end is unknown. We have argued that this limitation is paradoxically informative — it provides a plausible mechanistic explanation for why heat-treated preparations show stronger dose-response correlations — but this interpretation remains a hypothesis pending direct experimental verification through endpoint bacterial enumeration.

Third, strain taxonomic identities are not disclosed in this report, precluding species-level interpretation, genomic correlation, and identification of genetic determinants underlying the observed phenotypic differences. Full taxonomic characterization will be disclosed in subsequent publications. Fourth, substantial out-of-range values (up to 33.9% for MIP-1β above upper limit, 27.8% for IL-7 below detection limit) required imputation, introducing estimation uncertainty particularly at the extremes of the response range; the imputation methods used replaced out-of-range values with the empirically determined quantification boundaries of the assay-specific standard curve, as described in Section 2.6.1. The potential impact of imputed values on multivariate analyses was assessed by verifying that PCA component loadings, cluster assignments, and ML feature importance rankings were qualitatively stable when MIP-1β and IL-7 were excluded from the analysis matrix. The principal multivariate conclusions reported below are not dependent on the imputed values for these two analytes. Sensitivity analyses excluding MIP-1β and IL-7 from PCA, clustering, and ML analyses confirmed that principal component loadings, cluster assignments, and feature importance rankings were qualitatively unchanged. Formal quantitative reporting of these sensitivity analyses is deferred to future work with extended concentration ranges that would reduce the proportion of out-of-range values. Fifth, the three-point concentration range precludes formal pharmacological curve fitting (EC50, Hill coefficient); extension to 10⁴ and 10⁸ cells/well would complete the dose-response envelope and enable parametric modelling. Sixth, the 24-hour single timepoint captures peak IL-10, IL-12, and IFN-γ production (Kekkonen et al., 2008) but not the temporal dynamics of response initiation or attenuation. Seventh, the PBMC model lacks the epithelial barrier context noted by Kleiveland (2015); the Foligné IL-10/IL-12 correlation was established specifically for Gram-positive bacteria and does not extend to Gram-negative species; in vivo validation remains a necessary step. Notably, parallel evaluation of these same strains in an epithelial barrier model revealed fundamentally different response architectures: barrier function was viability-independent (best classification AUC = 0.662) in stark contrast to the viability-dependent cytokine responses reported here (AUC = 0.702), indicating that immunomodulatory and barrier-protective mechanisms have distinct molecular requirements — a dissociation that underscores the necessity of multi-endpoint characterization and will be reported in detail elsewhere. Eighth, the cross-validation approach for machine learning, while appropriate for this dataset size, provides an optimistic estimate of generalization performance compared to validation on an independent external dataset; the ML framework requires validation on independent PBMC datasets per the benchmarking standards of Li P. et al. (2025) and Zhou et al. (2020). Ninth, no positive control stimulus — such as lipopolysaccharide for monocyte TLR4 activation or phytohemagglutinin for T-cell activation — was included in the experimental design. The absence of a positive control means that inter-run PBMC functional competence cannot be formally verified from the reported data. While the robust cytokine responses observed across the majority of viable-bacteria conditions are themselves evidence of functional PBMC responsiveness, conditions showing low responses — particularly heat-treated Strain C and Strain I at 10⁵ TFU/well — cannot be distinguished with certainty from a globally suppressed PBMC preparation on that experimental day. Future studies employing this framework should include at minimum one positive control stimulus per experimental run.

### 4.8 Towards an Evidence-Based Dose Characterization Framework

Three operational principles emerge from the totality of findings. First, multi-concentration testing is necessary to characterize dose-response shape: monotonic scaling cannot be assumed without experimental verification, and bell-shaped kinetics are a mechanistically plausible, experimentally reproducible outcome for a non-negligible subset of strains. Second, viable and heat-treated preparations require independent characterization at matched concentrations: the Functional Equivalence Dose (FED_70_), grounded in biological and regulatory rationale (EMEA, 2010), provides a strain-specific benchmark for postbiotic dose justification that CFU-based approaches cannot supply. Third, machine learning-based functional classification of probiotic strains by cytokine-induction phenotype provides a data-driven annotation layer complementing the genomic approaches of Orkkatteri Krishnan et al. (2025) and Sun et al. (2022) and enabling evidence-based strain selection. Together, these principles constitute a pre-clinical characterization framework that could directly address EFSA’s dose-response evidence requirement (EFSA, 2012) and the ICH E4 mandate — providing the functional evidence base that regulatory health claim substantiation has consistently identified as absent.

The practical framework that emerges is straightforward: flow cytometry-enumerated bacteria at defined concentrations, 24-hour PBMC co-culture, multiplex cytokine quantification, and machine learning classification constitute a comprehensive immunomodulatory characterization pipeline completable within approximately 48 hours from bacterial preparation to functional annotation — a timeline compatible with industrial screening and regulatory dossier preparation.

Future studies should build on this framework in five directions. First, multi-donor validation across at minimum five donors of diverse sex, age, and immune backgrounds to establish population-level robustness of the dose-response profiles, cluster assignments, and ML classification accuracy. Second, whole-genome sequencing of the tested strains to enable genotype-phenotype correlation analysis, extending the Random Forest gene-trait matching approach of Meijerink et al. (2010) to the full multi-strain, multi-concentration dataset generated here. Third, time-course experiments capturing cytokine kinetics at multiple intervals (4, 8, 12, 24, 48 hours) to determine whether bell-shaped dose-response parameters are timepoint-dependent. Fourth, intestinal barrier model integration — incorporating epithelial co-cultures or immune-competent organoid systems alongside PBMC — would provide the physiologically complete assessment that the current model cannot offer, though such platforms remain technically immature for screening applications (Bar-Ephraim et al., 2020; Puschhof et al., 2021). Fifth, and most critically, in vivo validation of whether the immunophenotypic classification generated by this framework predicts clinical outcomes is the ultimate test of any preclinical screening tool. Only such validation can confirm whether the strain-specific pro-inflammatory signatures, anti-inflammatory profiles, or FED.

## 5. Conclusions

This study establishes a reproducible methodological framework that answers four questions fundamental to probiotic and postbiotic research. Are there differences between live and heat-treated bacteria? Yes — profound, consistent, and universal across the cytokine panel, with fold-changes spanning from 2.9-fold (IL-13) to 356.9-fold (TNF-α) across 16 of 17 analytes. Can dose-response relationships be demonstrated? Yes — particularly clearly in heat-treated preparations, where the absence of bacterial proliferation allows faithful observation of concentration-dependent immune activation (Spearman ρ up to 1.00), and in viable preparations for 6 of 10 strains across the tested 100-fold range. How should bacterial doses be measured? By flow cytometry per ISO 19344:2015, which provides the accuracy (RSD 6.70% vs 12.52% for plate counting), precision, and viability-state independence essential for meaningful dose-response characterization and live versus heat-treated comparison. Can machine learning enhance the analysis of well-structured experimental data? Yes — not only through high-accuracy viability classification (RF AUC-ROC 0.702)but through the emergence of three functional immunomodulatory phenotypes not part of the original experimental hypothesis, and through the identification of IL-12p70, IL-13, and IFN-γ, not IL-6 or IL-1β, as the primary discriminators of strain identity. These contributions establish a hypothesis-generating methodological and conceptual framework for evidence-based dose characterization, postbiotic functional equivalence assessment, and data-driven strain annotation, grounded in a single-donor model and requiring multi-donor validation before population-level conclusions can be drawn in probiotic and postbiotic research and product development. The strongly viability-dependent profile of Strain C — active when viable but attenuated after heat treatment — should be interpreted in the context of the absence of a positive control stimulus in this study; while robust responses in viable-bacteria conditions confirm general PBMC responsiveness, formal verification of maximal stimulatory capacity on the experimental day was not performed. All findings derive from a single-donor experimental model and constitute hypothesis-generating evidence requiring multi-donor validation before population-level conclusions can be drawn.

## Data Availability Statement

The datasets generated and analysed during the current study are available from the corresponding author on reasonable request.

## Funding

This study was conducted within the internal research and development activities of Probiotical Research Srl (Novara, Italy). No external or public funding was received.

## Author Contribution

M.P and A.A conceptualized and conducted the study. A.V. and G.D performed PBMC experiments. M.P and G.D performed data analysis. A.A. and M.P. Supervised. All authors reviewed and approved the manuscript.

## Acknowledgment

The authors acknowledge the use of Claude (Anthropic, claude.ai), a generative artificial intelligence assistant, in the preparation of the manuscript text, figure generation code, and machine learning analysis scripts. All AI-generated content was critically reviewed, verified against primary data sources, and edited by the authors. All interpretations, conclusions, and scientific claims are the sole responsibility of the authors.

## Conflict of Interest

All authors are employees of Probiotical Research Srl (Novara, Italy). Probiotical Research Srl is a subsidiary of Probiotical S.p.A which owns and commercially distributes probiotic strains, including those evaluated in the present study under anonymous strain codes (A–L). The authors declare that this employment relationship represents a potential conflict of interest, which has been disclosed in accordance with the journal’s editorial policy. The study design, data analysis, and interpretation were conducted independently of commercial considerations.

## Contribution to The Field

The probiotic field faces a structural paradox: products are commercialized at doses spanning three orders of magnitude without any functional evidence anchoring a specific concentration to a specific immunological outcome, and existing colony-forming unit (CFU)-based quantification methodology is inherently inapplicable to the growing category of heat-treated postbiotic preparations. This study addresses both gaps simultaneously and introduces a third layer of novelty, machine learning-based functional classification of probiotic strains by their cytokine-induction phenotype, for which no published precedent exists in the literature. By systematically characterizing ten Gram-positive probiotic strains in both viable and heat-treated forms across three concentration levels (10^5^, 10^6^, 10^7^ cells/well) using flow cytometric input standardization (Active Fluorescent Units/Total Fluorescent Units per ISO 19344:2015) and a 17-plex BioPlex cytokine readout in primary human PBMC from a single healthy donor, this work provides the first systematic multi-strain dose-response immunophenotyping dataset for Gram-positive probiotic strains. Three distinct immunological phenotype clusters are identified, including non-monotonic bell-shaped kinetics in two strains and strongly attenuated immunomodulatory activity in heat-treated form of one strain, establishing that dose-response shape and thermal treatment sensitivity are strain-intrinsic properties that cannot be generalized across species or formulations. Data-driven hierarchical clustering of cytokine fingerprints identified three candidate functional phenotype groups, internally re-recovered by Random Forest with 83.3% Leave-One-Out consistency — an internal partition-recovery metric that requires external validation on independent strain cohorts before the groups can be treated as generalizable biological phenotypes.

Seventeen cytokines were quantified simultaneously via Bio-Plex multiplex immunoassay. Live bacteria induced significantly greater responses across 16 of 17 cytokines, with tumor necrosis factor-alpha showing a 356.9-fold increase compared to heat-tretaed preparations. However, heat-treated bacteria exhibited stronger dose-dependency (Spearman rho up to 1.00), suggesting a more faithful representation of true dose-response relationships unconfounded by bacterial proliferation during co-culture.

Strain-specific profiling revealed three distinct immunomodulatory phenotypes emerging entirely from data-driven analysis without prior functional annotation.

## Notes

### Summary of Updates

This version updates the manuscript as follows: Terminology: throughout the text, figures, and supplementary material, "heat-inactivated" has been replaced with "heat-treated" and the abbreviation "HI" with "HT" when referring to the thermally treated preparations, to more accurately describe the experimental condition. Standard reagent terminology (heat-inactivated fetal bovine serum) and cited reference titles were left unchanged. Strain C narrative: descriptions of Strain C have been reframed from absolute terms (previously "inert" / "non-responsive") to a viability-dependent framing ("strongly viability-dependent", "attenuated activity following heat treatment"), which is both more accurate given the absence of a positive-control stimulus and less categorical. Section and figure headings were updated accordingly. Quantitative reconciliation: all reported values in the text, figures, and captions were systematically reconciled against the canonical master dataset to ensure full traceability, and minor discrepancies arising from rounding and transcription were corrected. Specific reconciliations included the feature-importance ranking (MCP-1 and IFN-gamma both 0.090), a co-regulation coefficient (IL-4/IL-5 corrected to 0.95), and the phrasing of significance counts; an unsupported per-strain F1 statement was removed. No underlying data were altered. Content organization: to meet the journal word limit, several sections (Cell Model rationale, Data Quality, and per-strain dose-response detail) were migrated to the Supplementary Material as Sections S1 to S5, with cross-reference callouts retained in the main text. All main figures remain in the main text. Supplementary figures: two supplementary figures were added, a classifier comparison across four machine-learning algorithms and a per-cytokine fold-change distribution plot. Figure 1B: the pathway-activation radar figure was regenerated from the canonical dataset with a documented, reproducible method, and its terminology updated. No new experimental data were added; all changes are corrections, clarifications, and reorganization of existing content.

