## Supplementary Material for "Beyond the Billion: Dose-Response Immunophenotyping and Machine Learning Classification of Live versus Heat-Treated Gram-Positive Probiotic Strains in Human Peripheral Blood Mononuclear Cells"

**Section S1. Cell Model: Rationale, Validation, and Analytical Gaps**

The peripheral blood mononuclear cell co-culture model was selected deliberately as the immunological readout platform for this study because it uniquely satisfies both the translational and analytical requirements of the experimental design. PBMCs are primary human cells, not immortalized lines with aberrant signaling, such as THP-1 monocytic leukemia cells that exhibit constitutive NF-κB activation and altered Toll-like receptor expression relative to primary monocytes (Bosshartm and Heinzelmann, 2016; Tedesco et al., 2018), or Jurkat T-cell lymphoma cells that harbor PTEN and SHIP1 loss resulting in constitutively hyperactivated PI3K/Akt signaling fundamentally distorting T cell receptor-dependent cytokine production (Abraham and Weiss, 2004). Critically, PBMCs constitute a heterogeneous population comprising monocytes, T lymphocytes, B lymphocytes, natural killer cells, and dendritic cells in physiological proportions (Kleiveland, 2015; Maecker et al., 2012). The 17-plex cytokine fingerprint that forms the basis for machine learning classification in this study emerges from the functional cross-talk among these subpopulations: monocytes and dendritic cells produce TNF-α, IL-6, and IL-12 through Toll-like receptor 2 engagement (Trinchieri, 2003), T helper cells generate IL-4 and IFN-γ, and natural killer cells produce IFN-γ mediated by dendritic cell-derived IL-12 through a positive feedback loop that cannot occur in single-cell-type cultures (Vivier et al., 2011; Iwasaki and Medzhitov, 2015). No single cell line can generate this dimensionality (de Jager et al., 2003; Duffy et al., 2014), and without it, data-driven functional classification would lack the biological signal required for meaningful phenotype discrimination. Intestinal organoids are epithelial structures devoid of immune cells by design (Sato et al., 2009; Clevers, 2016; Bar-Ephraim et al., 2020) and therefore measure barrier function rather than the integrated immune response; immune-competent organoid co-cultures remain technically immature, low-throughput, and unstandardized (Dijkstra et al., 2018; Neal et al., 2018), while organ-on-a-chip systems are fundamentally incompatible with the high-throughput factorial screening format required here (Ingber, 2022).

The PBMC model is, moreover, the only *in vitro* system for probiotic screening with documented clinical predictive validity. Foligné et al. (2007) demonstrated that PBMC-derived IL-10/IL-12 ratios predicted *in vivo* anti-inflammatory capacity in a murine TNBS-colitis model and that differences between blood donors did not prevent consistent strain ranking. This prediction was validated clinically by O'Mahony et al. (2005), confirmed at the human mucosal transcriptome level (van Baarlen et al., 2011), and translated into clinical trial strain selection in the PandA allergy prevention trial (Niers et al., 2005; Niers et al., 2009). No equivalent predictive validation chain exists for any cell line model, organoid system, or organ-on-a-chip platform in the context of probiotic immunomodulation (Papadimitriou et al., 2015; Klaenhammer et al., 2012).

The biological discriminatory power of the PBMC model across probiotic strains has been empirically quantified. Van Hemert et al. (2010) tested 42 *L. plantarum* strains with PBMCs from 3 donors and found IL-10 induction varied over a 14-fold range and IL-12 over a 16-fold range. The companion dendritic cell study by Meijerink et al. (2010) found even more dramatic variation: IL-10 ranged 39-fold and IL-12 ranged 600-fold across strains. Crucially, this study applied Random Forest gene-trait matching to identify six candidate genetic loci underlying the observed immunological differences. Notably, including the LamBDCA quorum sensing system and the plantaricin biosynthesis pathway, with gene deletion mutants confirming predictions, establishing the specific precedent for machine learning analysis of cytokine-induction data from probiotic-immune cell co-cultures that the present study extends to a multi-strain, multi-concentration, multi-preparation-type framework.

The substantial inter-donor variability in cytokine production capacity is genetically determined: Li et al. (2016) profiled PBMCs from 197 individuals and identified six genome-wide significant cytokine quantitative trait loci (QTLs), finding that cytokine production followed non-Gaussian or bimodal distributions with significantly increased inter-individual variability upon stimulation. Despite this variability, Foligné et al. (2007) and van Hemert et al. (2010) both demonstrated that relative strain rankings remain consistent across donors despite absolute value differences, validating the use of fold-change normalization and multiple donors as mitigation strategies. An important caveat noted by Kleiveland (2015) is that PBMCs "have a different composition, phenotype and activation status than cells found in intestinal tissue," lacking the epithelial barrier context and microbiome interactions present *in vivo*; the Foligné correlation, moreover, was established specifically for Gram-positive bacteria and should not be extrapolated to Gram-negative species. The PBMC model should therefore be understood as a validated comparative screening and ranking tool requiring complementary *in vivo* validation for translational conclusions.

Within the PBMC model specifically, dose-response data for probiotic immunomodulation remain strikingly scarce. Only three to four studies have attempted systematic multi-concentration characterization. Evrard et al. (2011) tested MOI 0.01–100 on dendritic cells and found strong dose-dependent DC maturation and cytokine induction with ~1,700 modulated genes at MOI 100 versus weak responses at MOI 0.01. Ghadimi et al. (2008) identified a dosage optimum of 10 bacteria per PBMC identical across all LAB tested, with bacterial genomic DNA contributing ~50% of effects dose-dependently. Hua et al. (2010) found that cytokine production decreased at $10^8$ CFU/mL compared to $10^6$–$10^7$ CFU/mL, providing early evidence for non-linear, inverted-U dose-response kinetics consistent with the hormesis concept. None of these studies applied the scale of the present work (10 strains × 2 preparation types × 3 doses × 17 cytokines), used preparation-type-neutral flow cytometric input standardization, or applied machine learning analysis to the resulting dataset. The machine learning gap in probiotic immunological research is equally pronounced. Standard ML algorithms are routinely applied to microbiome data for disease classification — Li P et al. (2025) benchmarked nine algorithms across 83 cohorts and 20 diseases, and Zhou et al. (2020) compared classifiers across 29 benchmark microbiome datasets. ML has been applied to cytokine profiles for clinical disease stratification: Wandong et al. (2022) demonstrated XGBoost classification of COVID-19 severity using IL-6 and IL-10 profiles with high discriminatory performance, establishing the precedent for ML on cytokine data in immunological contexts. Genomics-based probiotic classification has been developed by Sun et al. (2022), Orkkatteri Krishnan et al. (2025), and Shufang et al. (2024), achieving >95% accuracy for probiotic/non-probiotic binary classification from whole-genome features. Unsupervised ML has been applied to probiotic PBMC cytokine data by Coelho and Mossotto et al. (2020), who applied hierarchical clustering to cytokine profiles from PBMC stimulation assays targeting NOD2/TLR pathways, generating three distinct patient clusters — but their goal was patient stratification, not probiotic strain functional annotation. Wu et al. (2024) used unsupervised ML to differentiate 12 *Bifidobacterium* strains on an intestine-on-a-chip platform using organ-on-a-chip readouts rather than cytokine profiling. Charizani et al. (2024) described the first supervised ML model to predict *in vivo* probiotic potential from *in vitro* phenotypic features — but using cell surface properties (autoaggregation, hydrophobicity) in an aquatic/snail organism model, not cytokine profiles from human immune cells. Building on the ML-on-cytokine precedent established by Meijerink et al. (2010) in a single-species context, no published study has applied machine learning to classify probiotic strains by their immunomodulatory cytokine-induction fingerprint across a multi-strain, multi-dose, multi-preparation-type design in a primary human immune cell model.

The methodological weaknesses pervading the probiotic clinical trial literature compound all of the above gaps. Bafeta et al. (2018) found that among 384 probiotic RCTs, 28% reported no harms data, 80% did not report serious adverse events per group, and only 2% adequately reported all key safety components. McFarland et al. (2023) identified critical quality gaps across 206 probiotic systematic reviews, including inconsistent strain reporting and failure to account for dose differences. Suez et al.

(2019) highlighted that strain-level evidence is often inappropriately extrapolated, safety is under-evaluated, and personalized immune responses are routinely ignored. Szajewska et al. (2025) concluded that no data causally link probiotic clinical benefits to microbiota protection or recovery. Against this backdrop, the present study was designed to provide the first systematic multi-strain experimental characterization of dose-response relationships in probiotic immunology, using primary human PBMC from a single healthy male Caucasian donor, flow cytometric input standardization (AFU/TFU per ISO 19344:2015) applicable to both preparation types, 17-plex cytokine profiling, and machine learning-based functional strain classification. Ten Gram-positive probiotic strains were tested in both viable (AFU) and heat-treated (TFU) forms across three dose levels ($10^5$, $10^6$, $10^7$ cells/well), with three technical replicates per condition. Primary aims: (i) to determine whether systematic dose-response relationships exist across the tested concentration range, characterizing their shape, magnitude, and strain-specificity; (ii) to establish, using concentration-matched AFU/TFU comparison, the dose at which heat-treated preparations achieve functional equivalence to viable preparations, operationalized as the Functional Equivalence Dose (FED[70]); and (iii) to determine whether machine learning can classify probiotic strains into functionally meaningful immunological phenotype groups based on their 17-plex cytokine-induction fingerprint.

**Section S2. Data Quality and Distribution Characteristics**

The final dataset comprised 192 observations across 17 cytokines (90 viable conditions + 90 heat-treated conditions + 12 negative controls). All 10 probiotic strains were represented equally (18 samples each), with balanced distribution across the three bacterial concentrations (60 samples per concentration level). The balanced factorial structure ensured equivalent statistical power for all comparisons, with each cell of the design containing exactly three technical replicates.

Substantial out-of-range values were observed for several cytokines, reflecting the wide dynamic range of immunological responses inherent to the experimental design. MIP-1β exhibited the highest proportion of values exceeding the upper quantification limit (33.9%), indicative of potent chemokine induction particularly by viable bacteria. IL-7 showed the highest proportion below detection (27.8%), consistent with constitutively low expression in this experimental system. Raw cytokine concentrations spanned several orders of magnitude and exhibited pronounced right-skewness characteristic of immunological data. Log$^{10}$-transformation substantially improved distributional properties, with 16 of 17 cytokines showing increased Shapiro-Wilk p-values post-transformation. High coefficients of variation (25.8% for MCP-1 to 588.0% for IL-1β) reflected not merely technical noise but the substantial biological variability introduced by the 60-condition factorial design spanning two viability states and two orders of magnitude in bacterial concentration.

The overall pattern of immunomodulatory activity, visualised as mean log$^2$ fold-change relative to the unstimulated control (Figure 1), revealed heterogeneity operating at three levels simultaneously: response magnitude varies markedly across strains; the concentration-output relationship differs systematically between strains and between viable and heat-treated forms; and the cytokine-level composition of the immunological response shifts as a function of both dose and preparation type. Three analytes — IL-8, MIP-1β, and TNF-α — reached or exceeded the ULOQ of the BioPlex assay in a subset of conditions and are fully annotated in Supplementary Figure 1. To assess the impact of imputed values on multivariate analyses, key results — including PCA component loadings, cluster assignments, and ML feature importance rankings — were verified to be qualitatively stable when MIP-1β and IL-7 were excluded from the analysis matrix. The principal conclusions reported below are therefore not dependent on the imputed values for these two analytes.

**Section S3. Dose-Response Profiles of Viable Preparations**

Six of the ten viable preparations exhibited dose-dependent cytokine increases consistent across multiple analytes, establishing dose-response as a generalizable principle of probiotic-PBMC interaction (Figure 5). These data provide, at the 10-strain panel scale, the systematic multi-concentration evidence base whose absence was documented by Ouwehand (2017), Shane et al. (2010), and EFSA (2012). The strongest relationship was observed for Strain B: IL-6 increased from 4,892-fold (±SD) at $10^5$ AFU/well to 17,455-fold at $10^6$ and 18,081-fold at $10^{7:}$ IL-1β increased from 1,806-fold at $10^5$ to 8,829-fold at $10^6$ and 58,409-fold at $10^7$ — a 32-fold amplification across the full two-log concentration range. Strain A displayed concordant monotonic escalation: IL-6 from 1,410-fold at $10^5$ to 3,614-fold at $10^6$ and 13,548-fold at $10^7$; G-CSF from 33-fold to 217-fold to 7,954-fold; IL-10 from 103-fold to 549-fold to 7,028-fold. Strains F, H, I, and L similarly exhibited monotonic profiles (≥85% of cytokines showing V5 ≤ V6 ≤ V7) with varying magnitudes. Strain G showed a predominantly monotonic pattern (71% of cytokines) with minor deviations in IL-6 and IL-10 at the highest concentration. The existence of a minimum effective concentration and an optimal dose range where response scaling is steepest are directly apparent, providing the pre-clinical dose-characterization evidence that regulatory bodies require and that the field has been unable to supply.

**Section S4. Non-Monotonic Dose-Response Profiles: Bell-Shaped Kinetics**

Two strains — D and E — exhibited non-monotonic, bell-shaped kinetics with peak activity at $10^6$ AFU/well and measurable, multi-cytokine attenuation at $10^7$, consistent across technical replicates and consistent with the inverted-U dose kinetics reported by Hua et al. (2010) and the dosage optimum identified by Ghadimi et al. (2008). For Strain E: IL-6 increased from 1,379-fold at $10^5$ to a peak of 2,383-fold at $10^6$ before declining to 1,778-fold at $10^7$ (−25%); G-CSF showed the most pronounced pattern (1,055 → 1,480 → 246-fold, −83% from peak); IL-10 followed concordantly (308 → 377 → 95-fold). Strain D displayed a structurally parallel pattern across IL-6 (490 → 1,185 → 598-fold), G-CSF (61 → 411 → 263-fold), IL-10 (35 → 72 → 49-fold), with the peak uniformly at $10^6$. IFN-γ showed an inverse pattern for both strains, declining monotonically (Strain E: 52 → 44 → 24-fold; Strain D: 26 → 17 → 6-fold), indicating cytokine-specific rather than uniform attenuation. A critical dissociation was observed: heat-treated preparations for both strains showed stable or continuing activity at $10^7$ TFU/well for several analytes, arguing against non-specific cytotoxicity and in favour of an active-cell-dependent mechanism. The mechanistic interpretation of this dissociation is discussed in Section 4.

**Section S5. Strain C: A Strongly Viability-Dependent Immunomodulatory Profile**

Strain C occupied a singular position in the dataset. The viable form induced modest responses with cytokine-specific dose patterns: IL-6 showed no directional trend (1,142-, 800-, 1,014-fold across three dose points), while IL-10 displayed a progressive increase (21-, 57-, 180-fold). Overall immunostimulatory activity was substantially lower than high-magnitude strains such as B and A. The heat-treated form showed minimal induction across the cytokine panel at all concentrations: IL-6 (0.7, 0.9, 2.7-fold), IL-10 (0.2, 0.1, 0.4-fold), IL-1β (0.3, 0.3, 0.6-fold), G-CSF (<0.1-fold). At $10^7$ TFU/well — where heat-treated Strains B, H, and E elicit thousands-fold inductions — Strain C heat-treated preparations remained near baseline for 12 of 14 quantifiable analytes. This establishes that the strong attenuation of activity following heat treatment is a strain-specific biological property, not a generic consequence of microbial material at a given concentration, directly demonstrating that postbiotic activity cannot be assumed without explicit functional characterization of the specific heat-treated preparation.

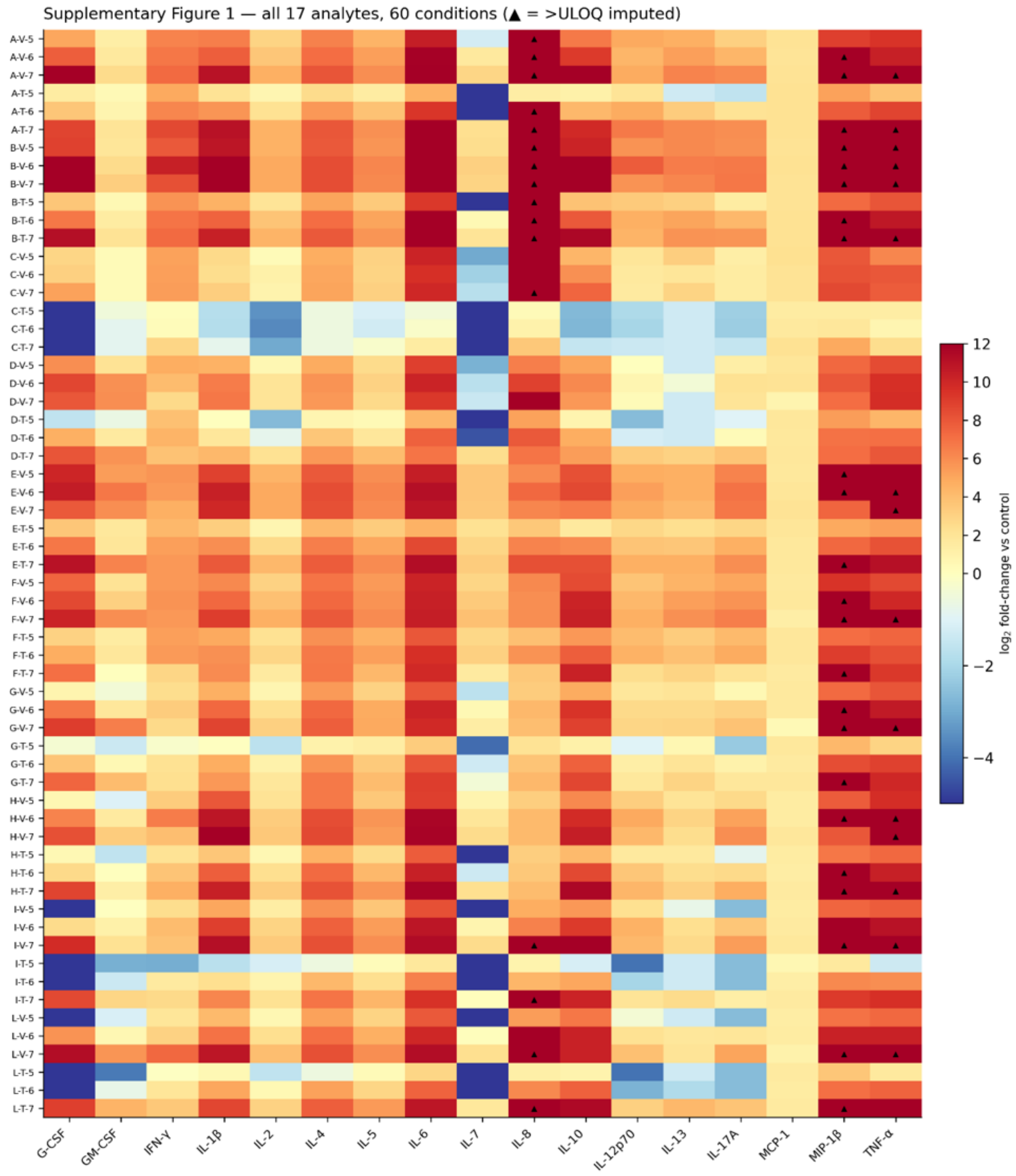


**Supplementary Figure 1.** Complete 17-analyte cytokine dataset. All 17 BioPlex analytes including IL-8, MIP-1β, and TNF-α. Black cells with white text ('>UL'): values exceeding the upper limit of quantification (IL-8 ULOQ ≈ $1.97\times10^{10}$ pg/mL; MIP-1β ULOQ ≈ $2.68\times10^{11}$ pg/mL; TNF-α ULOQ ≈ $6.24\times10^{10}$ pg/mL), present in 11, 18, and 14 of 60 conditions respectively. Row organisation as Figure 1: strains A–L, each with viable (V5, V6, V7) followed by heat-treated (HT5, HT6, HT7) preparations. Colour scale: diverging RdYlBu, capped at $\log_2$ = −5 (lower) and +12 (upper). (ULOQ values expressed as absolute concentrations in pg/mL as reported by Bio-Plex Manager software; fold-change values derived from these imputed concentrations are correspondingly extreme and are excluded from primary heatmap analyses)

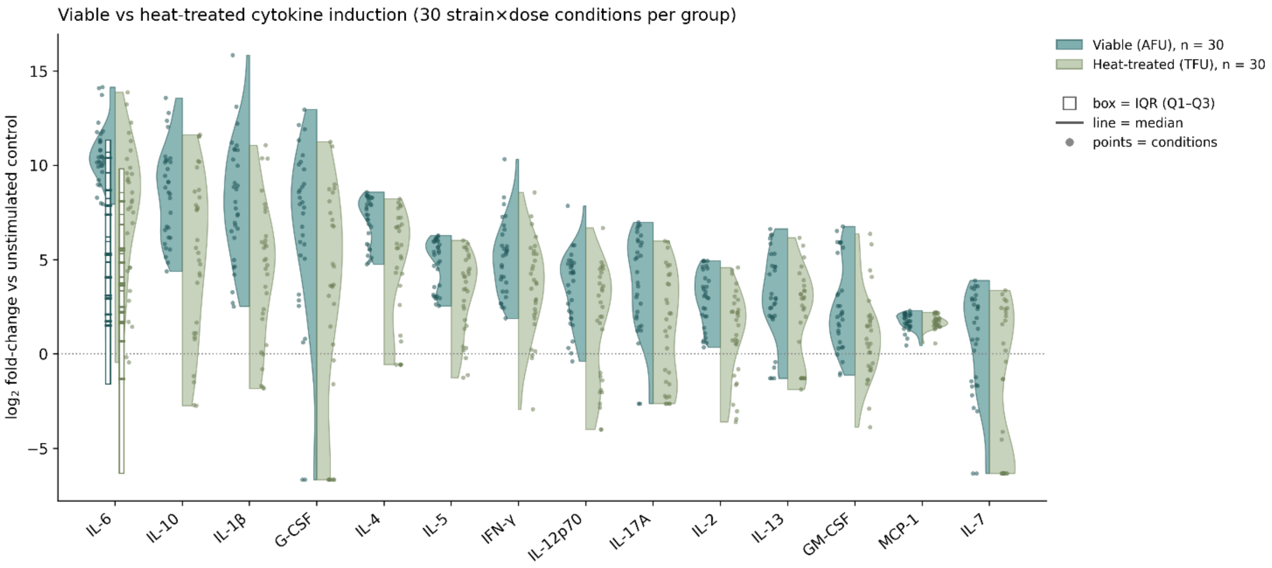


**Supplementary Figure 2.** Distribution of $\log_2$ fold-change (versus unstimulated control) for viable (AFU) versus heat-treated (TFU) preparations across 30 strain×dose conditions per group, shown per cytokine. Split violins show the full distribution; box = IQR (Q1–Q3), horizontal line = median, points = individual conditions.

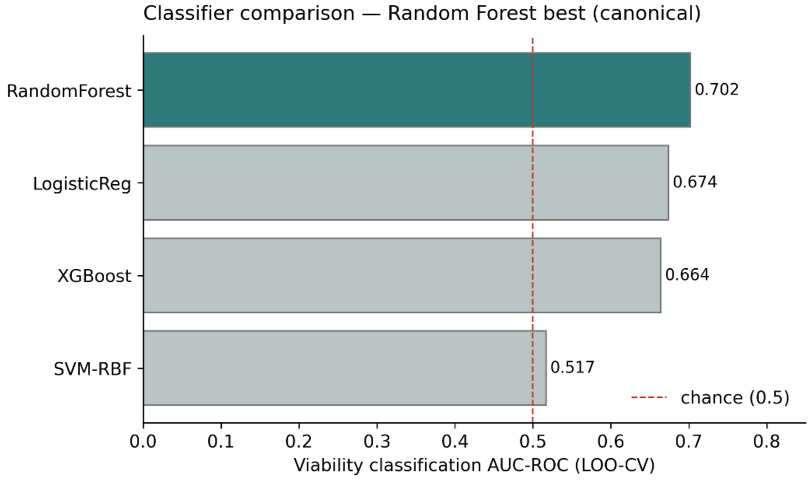


**Supplementary Figure 3.** Viability-classification performance across four machine-learning algorithms (leave-one-out cross-validation AUC-ROC). Random Forest achieved the highest performance (AUC 0.702); SVM-RBF performed near chance (0.517). Dashed line: chance level (0.5).
